# Time of arrival during plant disease progression and humidity additively influence *Salmonella enterica* colonization of lettuce

**DOI:** 10.1101/2024.02.16.580743

**Authors:** Megan H. Dixon, Dharshita Nellore, Sonia C. Zaacks, Jeri D. Barak

**Affiliations:** Department of Plant Pathology, University of Wisconsin, Madison, Wisconsin, USA; Microbiology Doctoral Training Program, University of Wisconsin, Madison, Wisconsin, USA

## Abstract

The interplay between plant host, phytopathogenic bacteria, and enteric human pathogens in the phyllosphere have consequences for human health. *Salmonella enterica* has been known to take advantage of phytobacterial infection to increase its success on plants, but there is little knowledge of additional factors that may influence the relationship between enteric pathogen and plant disease. In this study, we investigated the role of humidity and the extent of plant disease progression on *S. enterica* colonization of plants. We found that high humidity was necessary for replication of *S. enterica* on diseased lettuce, but not required for *S. enterica* ingress into the UV-protected apoplast. Additionally, the *Xanthomonas hortorum* pv. vitians (hereafter, *X. vitians*) *-* infected lettuce host was found to be a relatively hostile environment for *S. enterica* when it arrived prior to the development of watersoaking or following necrosis onset, supporting the existence of an ideal window during *X. vitians* infection progress that maximizes *S. enterica* survival*. In vitro* growth studies in sucrose media suggest that *X. vitians* may allow *S. enterica* to benefit from cross-feeding during plant infection. Overall, this study emphasizes the role of phytobacterial disease as a driver of *S. enterica* success in the phyllosphere, demonstrates how time of arrival during disease progress can influence *S. enterica’s* fate in the apoplast, and highlights the potential for humidity to transform an infected apoplast into a growth-promoting environment for bacterial colonizers.

**Importance:** Bacterial leaf spot of lettuce caused by *X. vitians* is a common threat to leafy green production. The global impact caused by phytopathogens, including *X. vitians*, is likely to increase with climate change. We found that even under a scenario where increased humidity did not enhance plant disease, high humidity had a substantial effect on facilitating *S. enterica* growth on *Xanthomonas-*infected plants. High humidity climates may directly contribute to the survival of human enteric pathogens in crop fields or indirectly affect bacterial survival via changes to the phyllosphere brought on by phytopathogen disease.

## INTRODUCTION

Agricultural fields serve as habitats for diverse communities of microorganisms, including bacteria. Although the surface of a leaf may seem innocuous, bacteria that colonize the phyllosphere must adapt to hostile conditions including nutrient scarcity, extreme fluctuations in moisture availability, UV irradiation, and the host immune response (1, 2). To increase survival, some bacteria abandon the leaf surface and enter the apoplast where they may evade such stresses, so long as they can evade or suppress the plant immune response. Phytobacterial pathogens overpower host defenses and destructively (3–8). For example, *Xanthomonas hortorum* pv. vitians (hereafter, *X. vitians*) causes bacterial leaf spot of lettuce(9, 10), which is characterized by symptom development of dark, water-soaked leaf lesions (9, 10)(11). Phytobacterial disease causes yield loss (7, 12–18)(11), which devastates agricultural industries worldwide and is already a major cause for concern on its own. However, growing evidence of *Salmonella enterica’s* potential to benefit from host changes brought on by phytopathogen infection (7, 12–18) supports that phytobacterial disease may be linked to another issue: food-borne illness.

*Salmonella enterica* is an environmental microbe that effectively colonizes both mammalian and environmental hosts (19), relying on its ability to adapt and survive in diverse niches (20). Irrigation water, soils, and plant surfaces are all examples of places where *S. enterica* can persist for long periods of time, months to years (21–23). *S. enterica* is one of several enteric pathogens recognized as colonizers of plants (24), and salmonellosis outbreaks have been linked to consumption of raw produce (25). Given the prevalence of phytobacterial disease on agricultural crops, *S. enterica* is likely to interact with pathogenic phytobacteria when *S. enterica* is present in a field. Considering the public health threat posed by *S. enterica* colonization of crop plants, studying biotic and abiotic factors that influence its ability to survive in the environment is a critical area of research.

Phytobacterial disease promotes the survival of *S. enterica* on plants pre-harvest (7, 12–18). A recently published study from our lab showed that tomato leaves infected by *X. hortorum* pv. gardneri enabled *S. enterica* cells that arrive on the leaf surface to become internalized within the leaf apoplast and replicate (17). After completing this work, we questioned whether *S. enterica* would benefit from any bacterial leaf spot disease. Lettuce has been implicated in food-borne illness outbreaks caused by *S. enterica*, and lettuce is susceptible to the bacterial spot disease pathogen, *X. vitians* (26–28). Because lettuce is commonly eaten raw, without food safety intervention strategies that can lower pathogen populations below the infectious dose, we determined that studying the romaine lettuce-*X. vitians* pathosystem would be key to expanding our knowledge of factors that influence *S. enterica* population size on plants.

The goal of our study was to examine how *X. vitians* disease and environmental factors, specifically high relative humidity, may impact the fate of *S. enterica* in the apoplast of lettuce plants. By studying another pathosystem, we can discover differences that may exist for *S. enterica* plant colonization driven by host. Additionally, we investigated how *S. enterica’s* success on *X. vitians*-infected lettuce could be influenced by how early or late during infection that *S. enterica* arrives, effectively using *S. enterica* as a biological reporter for host changes during phytobacterial infection and disease progress. Lastly, we measured how *S. enterica* growth in a sucrose-rich environment is affected by *X. vitians in vitro* to determine a mechanism by which *S. enterica* replicates in the presence of *X. vitians* infection.

## METHODS

### Bacterial strains, media, and culture conditions

Bacterial strains used in this study are listed in Table 1. *X. vitians* cultures were prepared by inoculating 3 mL of nutrient broth with frozen cells and incubating at 28°C with shaking at 200 rpm for 2 days. *Salmonella enterica* cultures were prepared in a similar manner but incubated in lysogeny broth (LB) overnight. Cultures were then diluted in sterile water and prepared to the desired cell concentration. Nutrient agar (NA) was used for enumeration of *X. vitians,* and LB agar amended with 20 μg/mL of nalidixic acid was used for enumerating *S. enterica*. Agar plates were incubated for ∼48 hrs at 28°C or 16-24 hr at 37°C for *X. vitians* and *S. enterica*, respectively.

**TABLE 1.** Bacterial strains used in this study.

| Strain name and description | Reference or source |
| --- | --- |
| *Xanthomonas hortorum* pv. *vitians* L7 (*X. vitians*) | J. Jones |
| *Xanthomonas hortorum* pv. *vitians* L7; Cm<sup>R</sup> transformed with pNO-mCherry | This study |
| *Escherichia coli*; Cm<sup>R</sup>; pNO-mCherry | Z. Gitai |
| *Salmonella enterica* serovar Typhimurium 14028s; Nal<sup>R</sup> | (82) |
| *Salmonella enterica* serovar Typhimurium 14028s; Kan<sup>R</sup>; transformed with pKT-Kan; GFP | (83) |

<sup>R</sup> = resistance to chloramphenicol (Cm), nalidixic acid (Nal), or kanamycin (Kan)

A fluorescently labeled *X. vitians* strain was produced for this study by transforming a wild-type *X. vitians* L7 strain with high-copy plasmid pNO-mCherry (gifted from Z. Gitai lab; modified version of plasmid described in (29)) derived from pTH18Cr (30), which carries a chloramphenicol resistance cassette and constitutively expresses mCherry via P_L*tet*O-1_ (31) promoter with *tet* repressor removed. Plasmid DNA was isolated from *Escherichia coli* using a Spin MiniPrep kit (Qiagen) and transferred to *X. vitians* via electroporation.

### Bacterial inoculation of lettuce

*Lactuca sativa* (romaine lettuce) seedlings were cultivated in growth chambers with a 16-h photoperiod at 24°C. Lettuce plants (4-5 weeks old) were infiltrated with *Xanthomonas vitians* inoculum prepared to an OD_600_ = 0.2 (∼3 x 10^8^ CFU/mL) or sterile deionized H_2_O as a control. *Xanthomonas* infiltration was performed via needleless syringe on the abaxial side of the leaf as described previously (32) and adapted for lettuce leaves. “Middle leaves” (the 3^rd^ or 4^th^ oldest true leaves) were selected for infiltration, which were fully open and relatively flat. The approximate volume infiltrated into each infiltration zone was 15 μL, which produced infiltration zones 1-1.5 cm long. For each treatment group, 6-8 plants were infiltrated per experimental replicate with 1-2 leaves infiltrated per plant. Biological replicates were considered the individual plants.

Infiltrated lettuce plants were maintained in large plastic bins with a 16-h photoperiod at ∼24°C and a moderate relative humidity (RH) of 50-70%. Lettuce plants were subsequently inoculated with *Salmonella enterica* cell suspensions at the given time post-infiltration with H_2_O or *X. vitians* cells. *S. enterica* was diluted in sterile deionized water to a cell concentration ∼10^6^ CFU/mL. *S. enterica* cell suspension droplets (3 uL; ∼3000 CFU/droplet) were pipetted onto the adaxial surface of each infiltration zone to mimic arrival via irrigation or splash dispersal, avoiding overlap between the droplet site and any microscopic damage caused by syringe infiltration. Leaves were then destructively sampled at given time points (1–5) post-*S. enterica* arrival. When experiments required sampling beyond 1-day post-*S. enterica* arrival, lids were placed on the bins at 24 hours post-*S. enterica* arrival to raise the RH to >90%. Plants were kept at a RH >90% unless noted; when lids were left off or removed, RH level was 55-70%.

### Bacterial population sampling

Lettuce leaves were cut from the plant and when applicable, UV irradiation was utilized to inactivate *S. enterica* cells colonizing the leaf surface and thus measure *S. enterica* colonizing the apoplast (32). Leaf tissue was irradiated with UV-C (254 nm) by placing samples adaxial side facing up inside a Stratalinker UV Crosslinker 1800 and dosing the tissue at an exposure level of 150,000 μJ/cm^2^. For the experiment described in **Figure 1**, for each plant, one of two inoculated middle leaves was randomly selected for UV irradiation and the other leaf left non-irradiated as a control. For all other *in planta* experiments involving UV irradiation treatment, middle leaves that had been inoculated with *S. enterica* cells on both sides of the midrib were split in half after sampling—one half of the leaf was randomly selected for UV irradiation and the half left non-irradiated. Similar to what was demonstrated previously with *S. enterica*-inoculated tomato leaves (32), UV irradiation treatment reduced the populations of *S. enterica* applied to the lettuce leaf surface by at least 2 logs. Preliminary experiments demonstrated that *S. enterica* cells infiltrated directly into the apoplast of lettuce leaves were not affected by UV irradiation treatment (data not shown).

**Figure 1:**
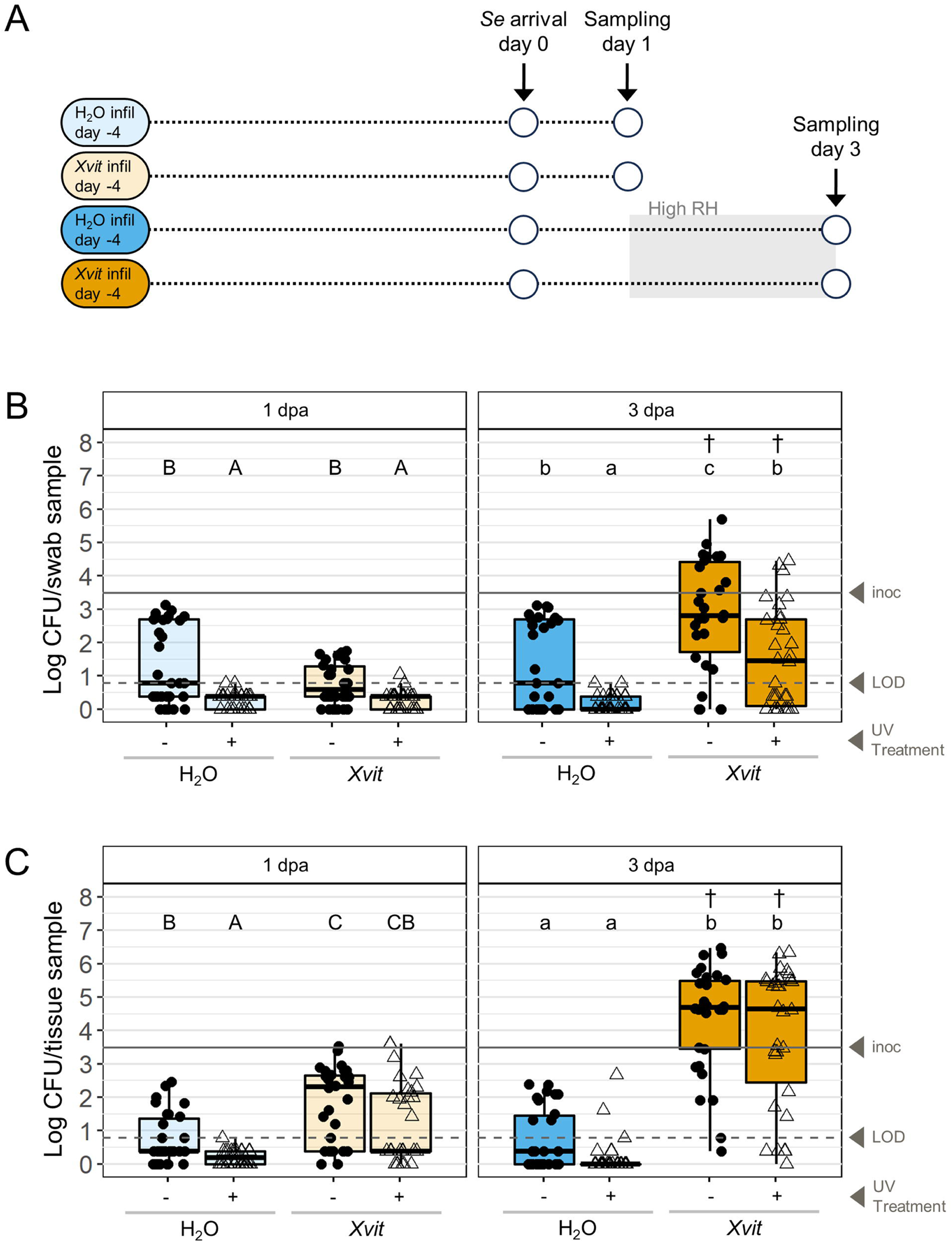
*X. vitians* infection facilitates *S. enterica* replication and colonization of UV-protected niches *in planta*. **(A)** Schematic of experimental design including color coded treatment descriptions. *S. enterica* (*Se*) cells were applied to leaves previously infiltrated with H_2_O or *X. vitians* (*Xvit*) at 4 dpi, and plants were sampled at 1-or 3-days post-*Se* arrival (dpa). High humidity (>90% RH) exposure is depicted as grey shading. **(B-C)** *S. enterica* (*Se*) colonization of H_2_O-infiltrated (blue boxplots) and *X. vitians*-infiltrated (orange boxplots) plants as measured from surface swab (**B**) and homogenized lettuce tissue **(C)** at 1-and 3-days post *Se* arrival (dpa). Darker boxplot colors for 3 dpa data signify exposure to high humidity prior to sampling. Leaves were irradiated with UV immediately prior to sampling (unfilled triangle points) or left non-irradiated (filled circle points). Letters above the boxplots indicate significance between treatment groups among leaves sampled at 1 dpa (uppercase) and among leaves sampled at 3 dpa (lowercase) based on Welch-ANOVA and *post hoc* Games-Howell means comparison tests (*P* < 0.05). Daggers (†) indicate significant increases in *Se* colonization between 1 dpa and 3 dpa within the same infiltration treatment and UV treatment groups based on two-sample T-tests (*P* < 0.05). Solid line indicates average *Se* arrival population applied to each sampled area, and dashed line indicates limit of detection (LOD). Values below LOD are defined in Materials and Methods. *N* = 30 leaves/treatment group; data is pooled from five independent experimental replicates.

For one experiment, measurement of surface *S. enterica* populations was determined by removing cells from sampled leaves using swabs dipped in Tween-20 as previously described (32). The washate from the swab was then plated to quantify viable surface cells. For all sampled leaves, bacterial colonization of leaf tissue was measured by excising 1-cm-diameter leaf disks from infiltration zones using a tissue corer and homogenizing the tissue in 500 uL of sterile water using a Dremel tool (33). Leaf homogenate was dilution-plated to enumerate *S. enterica* and *X. vitians* populations. The limit of detection (LOD) for *S. enterica* colonization was 5 CFU/leaf disk (or 10 CFU/sample for the summation of swab and leaf tissue *S. enterica* populations). The remaining leaf homogenate was enriched with LB containing nalidixic acid (20 μg/mL) to determine if *S. enterica* populations below the LOD were present in samples for which direct plating did not result in the growth of *S. enterica*. For non-detects, LB-enriched samples were plated ∼24 hours after enrichment. *S. enterica* colonization datapoints plotted at zero were confirmed to be zero based on the absence of *S. enterica* in the LB-enriched sample. Data plotted halfway between zero and the limit of detection indicates absence of colonies via direct plating and a positive enrichment sample.

### Electrolyte leakage measurements

Lettuce plants were prepared by infiltrating two adjacent middle leaves. The relatively older leaf was infiltrated four times, with two infiltration zones placed on each side of the leaf margin. One infiltration zone was randomly selected for bacterial population enumeration, and the remaining three infiltration zones were used for electrolyte leakage measurements. Electrolyte leakage was measured as performed previously (34, 35). In brief, for each biological replicate, three 1 cm diameter tissue disks were excised from a single leaf and pooled. Leaf tissue was placed in 12-well plates with wells containing 4 mL of water each. An EcTestr 11 (Eutech Instruments) conductivity tester was used to measure the leakage of electrolytes from the tissue into the water after 6 hours of incubation. Conductivity was reported by subtracting blank measurements taken immediately following a 45-minute wash step (time = 0 hours) from measurements taken at 6 hours.

### Confocal laser scanning microscopy

Lettuce plants were infiltrated with sterile deionized H_2_O or *X. vitians* L7 + pNO-mCherry cells prepared to an OD_600_ = 0.2 and incubated in plastic bins under moderate RH (50-70%) as described above. Droplets of *S. enterica* + pKT-Kan (GFP-labelled) cell suspensions containing ∼10^5^ CFU were applied to the surfaces of infiltrated zones at 4-days post-infiltration with H_2_O and 1-or 4-days post-infiltration with *X. vitians*. Inoculated plants were then incubated for 2 days, with high humidity (RH >90%) applied for the final 24 hours prior to sampling. Whole leaves were excised from the plants and kept in petri plates containing a moist Kim-wipe prior to imaging. Slides were prepared immediately before imaging by excising 0.5 cm diameter tissue disks at the site of *S. enterica* application and wet mounting them between two glass cover slips.

Confocal laser scanning microscopy (CLSM) images were obtained using a Zeiss Elyra LSM780 inverted confocal microscope and 40x water immersion 1.10 C-Apochromat objective. The 488 and 561 nm lines of an argon laser were used for excitation of GFP, mCherry, and chlorophyll A autofluorescence, which were detected at 500-550 nm (GFP), 552-624 (mCherry), and 630-700 nm (chlorophyll A) using the Zen (Zeiss) software. GFP and chlorophyll A autofluorescence were detected using Track 1, and mCherry detected using Track 2, with track switching set to every line. Master gain for GFP, mCherry, and chlorophyll A was set to 700, 750, and 850, respectively. A transmitted light detector (with master gain set to 450) was used for capturing differential interference contrast (DIC) images, which are shown overlaid with confocal fluorescence imaging data. For each treatment, leaf samples from 2-3 plants were imaged and 9-15 image fields were captured at both the surface level and within the apoplast at a z-depth = 4.6-14.5 μm (mean = 8.7 μm) relative to the surface (z = 0). Observations of the presence/absence of GFP-labelled *S. enterica* cells within the apoplast were recorded for every field.

For representative images shown, image processing via ImageJ was applied consistently across images, as performed previously (32). For DIC images, brightness and contrast were linearly adjusted using the “reset” button within the “Brightness/Contrast” tool. To improve visibility of *S. enterica* and *X. vitians* cells, the “enhance contrast” tool was applied to the GFP and mCherry channels with level of saturated pixels set to 0.05% and “equalize histogram” box left unchecked. Chlorophyll A channel data was omitted for representative images.

### Sucrose media co-culture assay

M9 minimal media containing sucrose or glucose as the sole carbon source (hereafter, M9 sucrose and M9 glucose) was prepared fresh for every experimental replicate with the following components diluted in de-ionized water: MgSO_4_ (2.0 mM), D-glucose or D-sucrose (0.4% w/v), CaCl_2_ (0.10 mM), Na_2_HPO_4_ (47.8 mM), KH_2_PO_4_ (22.0 mM), NaCl (8.6 mM), and NH_4_Cl (18.7 mM).

Stationary phase *X. vitians* cells grown in NB were washed three times in 1X phosphate-buffered saline (PBS) and normalized to an OD = 0.2. Normalized culture was inoculated into M9 sucrose and incubated at 28°C with shaking. Separate *X. vitians* culture flasks were incubated for either 24 or 48 hours in advance of *X. vitians* culture or cell-free supernatant (CFS) being introduced to *S. enterica.* CFS was prepared by successively filtering *X. vitians* culture through 0.8 μm and 0.2 μm filters. Sterility of CFS was confirmed via nutrient broth enrichment.

*S. enterica* cells grown overnight in LB were prepared by washing stationary phase cells three times in 1X PBS, normalizing to an OD = 0.1, and diluting 1000-fold in sterile water for a final concentration of ∼10^5^ CFU/ml. Diluted *S. enterica* cells or sterile water (100 μL) was then combined with 1.4 mL of *X. vitians* cell cultures or CFS per well of a 12-well plate. *S. enterica* and *X. vitians* cell concentrations were measured for triplicate wells immediately after 12-well plates were prepared at time = 0 hours and following 28 hours of incubation at 28°C.

### Statistical analysis

Statistical analyses and plot generation were performed using R Studio version 2023.03.1 (Posit team, RStudio: Integrated Development Environment for R. Posit Software, PBC, Boston, MA.[http://www.posit.co/]). Data from three independent replicate experiments were aggregated, unless otherwise noted in the figure legend. The number of biological replicates for each replicate experiment was 6-8 lettuce plants. For the *in vitro* growth assay in sucrose media, the number of experimental (biological) replicates was 4, with each biological replicate calculated from the mean of three technical replicates. Biological replicates for the *in vitro* growth assay are defined as independent sub-cultures of the frozen bacterial stocks whereas technical replicates refer to replicate wells of a 12-well plate. Bacterial population data were plotted following a log_10_(x+1) transformation. Statistical analysis was performed using log-transformed bacterial population data.

Welch’s analysis of variance (ANOVA) and post-hoc Games-Howell tests, which do not assume equal variance between treatment groups, were used for comparing *in planta* bacterial populations among treatment groups. When comparing bacterial populations between timepoints, a Welch two-sample T-test was used, and statistical differences displayed above boxplots using an asterisk symbol (*). When *S. enterica in planta* populations were compared to the amount of *S. enterica* applied, arrival population, a Wilcoxon one-sample test was used, and statistical differences indicated using a dagger symbol (†). Lastly, Fisher’s exact test was used for comparing proportions of *in planta* samples with *S. enterica* colonization that exceeds the arrival population. The significance threshold for all statistical comparisons was a *P* value of <0.05.

## RESULTS

### *X. vitians*-infected lettuce enables *S. enterica* growth and colonization of UV-protected niches

To determine whether bacterial spot of lettuce, which is caused by *X. vitians*, can facilitate *S. enterica* growth and colonization of UV-protected niches, lettuce plants were infiltrated with H_2_O or *X. vitians* cells and treated with *S. enterica* cells at 4-days post-infiltration (**Figure 1A**). The number(32) of *S. enterica* cells recovered by swab was significantly lower for UV-irradiated leaves compared to non-irradiated leaves at 1 dpa (**Figure 1B).** Furthermore, the number of *S. enterica* cells recoverable from the surface of the lettuce tissue by swab at 1-day post-*S. enterica* arrival (dpa) was not significantly different between healthy and *X. vitians*-infected plants (**Figure 1B**; *P* = 4.43 x 10^-7^, Welch ANOVA). However, the number of *S. enterica* cells recovered from the swab was significantly higher for *X. vitians-*infected leaves at 3 dpa compared to healthy leaves (**Figure 1B**; *P* = 9.17 x 10^-11^, Welch ANOVA) suggesting that *S. enterica* populations successfully colonized the leaf surface of infected leaves once disease progressed. For *X. vitians-*infected leaves, the *S. enterica* populations measured from the swab also increased between 1 and 3 dpa for both non-irradiated and UV-irradiated leaves, whereas the number of *S. enterica* cells recovered from the leaf surface did not increase at 3 dpa for healthy leaves (**Figure 1B**).

For the same leaves, *S. enterica* populations were quantified for the leaf tissue itself following the swabbing of the surface (**Figure 1C**). The *S. enterica* populations measured from these tissue samples represent cells that failed to be physically removed by swab—either due to attachment to the tissue or colonization of microscopic grooves or internal spaces. For UV-irradiated leaves in particular, the measured *S. enterica* populations represent cells colonizing UV-protected niches. We found that at 1 dpa, the number of *S. enterica* cells measured from the lettuce tissue was generally higher for *X. vitians-*infected lettuce than healthy lettuce, as demonstrated by the higher *S. enterica* populations measured on both non-irradiated and UV-irradiated leaves (**Figure 1C**; *P* = 6.79 x 10^-13^, Welch ANOVA). This contrasts with what was observed with the swab sampling, where *X. vitians* infection did not impact the number of *S. enterica* cells removable from the lettuce surface at the earlier sampling time, 1 dpa (**Figure 1B**). At 3 dpa, the number of *S. enterica* cells associated with the lettuce tissue was significantly higher on *X. vitians*-infected leaves compared to healthy leaves, and the *S. enterica* populations associated with *X. vitians-*infected tissue increased over time demonstrating *S. enterica* replication (**Figure** 1C; *P* = 8.36 x 10^-21^, Welch ANOVA).

### *S. enterica* growth on *X. vitians*-infected lettuce relies on high humidity

After establishing that *X. vitians* infection promotes *S. enterica* growth and colonization of UV-protected niches, we considered other factors that might influence *S. enterica’s* growth, such as humidity. We investigated whether the combination of *X. vitians* infection and high humidity (>90% RH) could additively benefit *S. enterica.* We tested the hypothesis that high humidity maximizes *S. enterica’s* ability to grow on *X. vitians*-infected plants by measuring *S. enterica’s* colonization of healthy and *X. vitians-*infected leaves at 4 days post-arrival (**Figure 2A**). We found that exposure to 72 hours of high humidity (Humidity Treatment III) resulted in greater *S. enterica* populations on *X. vitians-*infected plants compared to *S. enterica* populations on *X. vitians*-infected plants that did not experience high humidity (Humidity Treatment I, **Figure 2B**; *P* = 2.60 x 10^-18^, Welch ANOVA). In addition, a full 72 hours of exposure to high humidity resulted in greater *S. enterica* populations on *X. vitians*-infected lettuce compared to a fluctuating humidity treatment in which 24 hours of high humidity was followed by 48 hours of drier conditions (Humidity Treatment II). Furthermore, 72 hours of exposure to high humidity resulted in *S. enterica* growth, based on comparison of *in planta S. enterica* populations to the amount of *S. enterica* cells in the arrival population (**Figure 2B**). Meanwhile, 24 hours of exposure to high humidity did not promote *S. enterica* growth but supported *S. enterica* survival at a population level similar to the arriving population. For plants that were not exposed to high humidity at all (Humidity Treatment I), *X. vitians* infection resulted in greater *S. enterica* colonization compared to healthy leaves (**Figure 2B**), demonstrating that *X. vitians* infection influences *S. enterica* survival even in the absence of prolonged high humidity conditions. Contrastingly, *S. enterica* populations on healthy leaves were not significantly influenced by high humidity (*P* > 0.05), and *S. enterica* populations for all healthy plants regardless of humidity treatment were below the arrival population level at 4 days post-*S. enterica* arrival (**Figure 2B**). Additionally, fluctuating humidity (Humidity Treatment II) resulted in a reduction in *X. vitians* populations compared to constant moderate humidity conditions (**Supplementary** Figure 1; *P* = 0.036, Welch ANOVA). *X. vitians* and *S. enterica* populations were not found to be correlated (*P* > 0.05, Pearson correlation), demonstrating that *S. enterica’s* response to increasing exposure to high humidity on *X. vitians*-infected leaves is attributed directly to humidity and not *X. vitians* population density. In fact, high humidity introduced during early infection at 3 dpi was found to disrupt *X. vitians* disease progress, based on the observations that *X. vitians* populations and electrolyte leakage measurements were lower at 4 days post-*X. vitians* infiltration for lettuce plants exposed to 24 hours of high humidity (>90% RH) prior to sampling **(Supplementary** Figures 2A-C).

**Figure 2:**
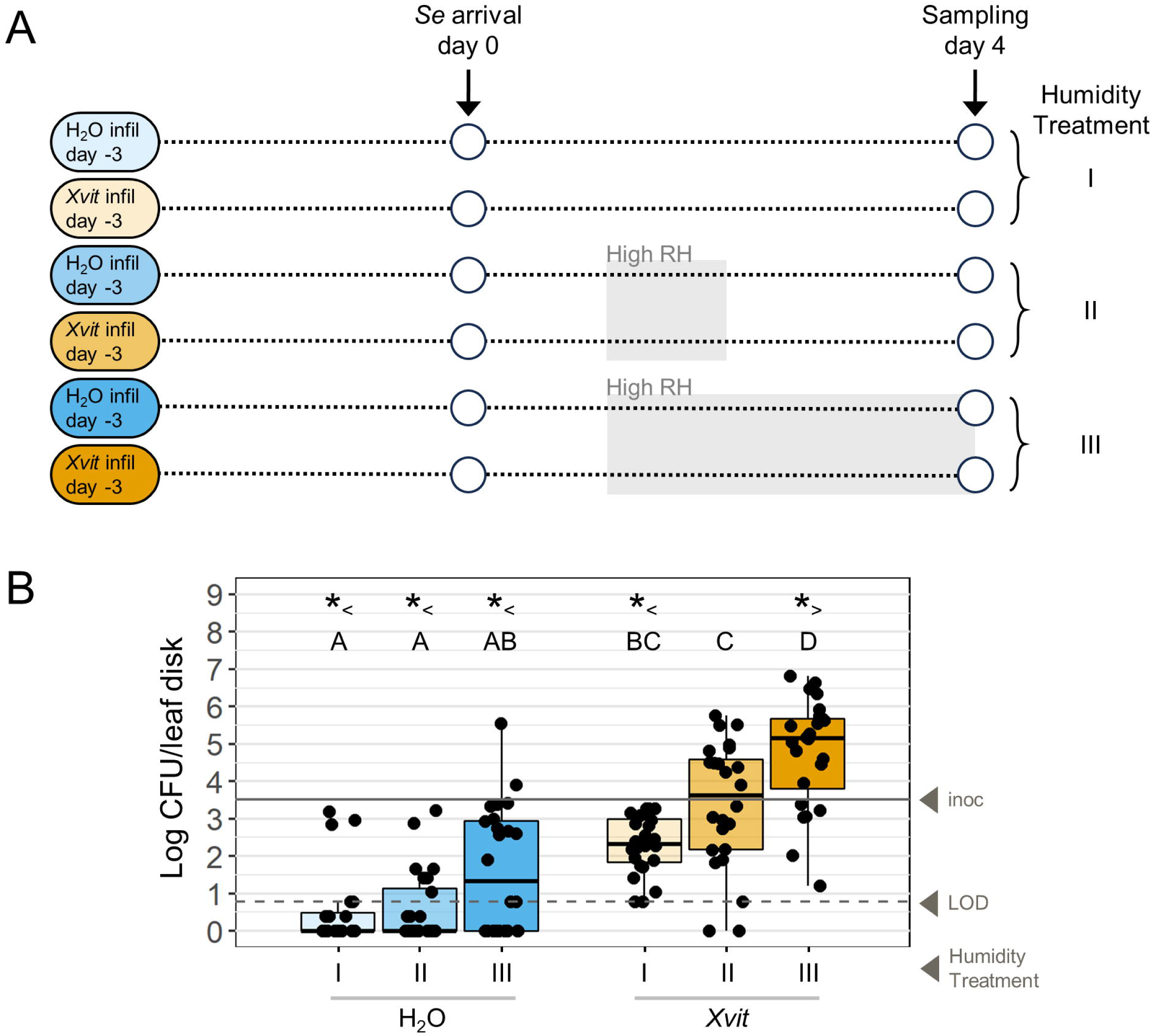
High humidity (>90% RH) exposure and *X. vitians* infection additively enhance *S. enterica* colonization of lettuce. **(A)** Schematic of experimental design. Lettuce plants were divided into six treatment groups. *S. enterica* (*Se*) cells were applied to leaves previously infiltrated with H_2_O or *X. vitians* (*Xvit*) at 3 days post-infiltration. Plants were sampled following 0, 24, or 72 hours of exposure to high humidity (>90% RH) at 4 days post-*Se* arrival (Humidity Treatments I, II, and III, respectively). **(B)** *Se* colonization of H_2_O-infiltrated (blue boxplots) and *X. vitians*-infiltrated (orange boxplots) plants at 2 days post-arrival (dpa) in response to varying humidity treatments. Letters above the boxplots indicate significance between treatment groups based on Welch-ANOVA and *post hoc* Games-Howell means comparison tests (*P* < 0.05). Asterisks indicate that *Se in planta* colonization is significantly less than (*<) or greater than (*>) the arrival population based on one-sample Wilcoxon tests (*P* < 0.05). Solid line indicates average *Se* arrival population applied to each sampled area, and dashed line indicates limit of detection (LOD). < LOD values are defined in Materials and Methods. *N* = 24 leaves/treatment group; data is pooled from three independent experimental replicates.

### High humidity is not required for *S. enterica* colonization of the apoplast of *X. vitians-*infected leaves

Having previously shown that water-congestion within the apoplast of tomato leaves is sufficient for *S. enterica* ingress into *Xanthomonas*-infected leaves (32), we hypothesized that humidity could influence *S. enterica* colonization of UV-protected niches of lettuce. To test this hypothesis, we applied *S. enterica* to lettuce plants infected with *X. vitians* and subjected the lettuce plants to different humidity treatments (**Figure 3A**).

**Figure 3:**
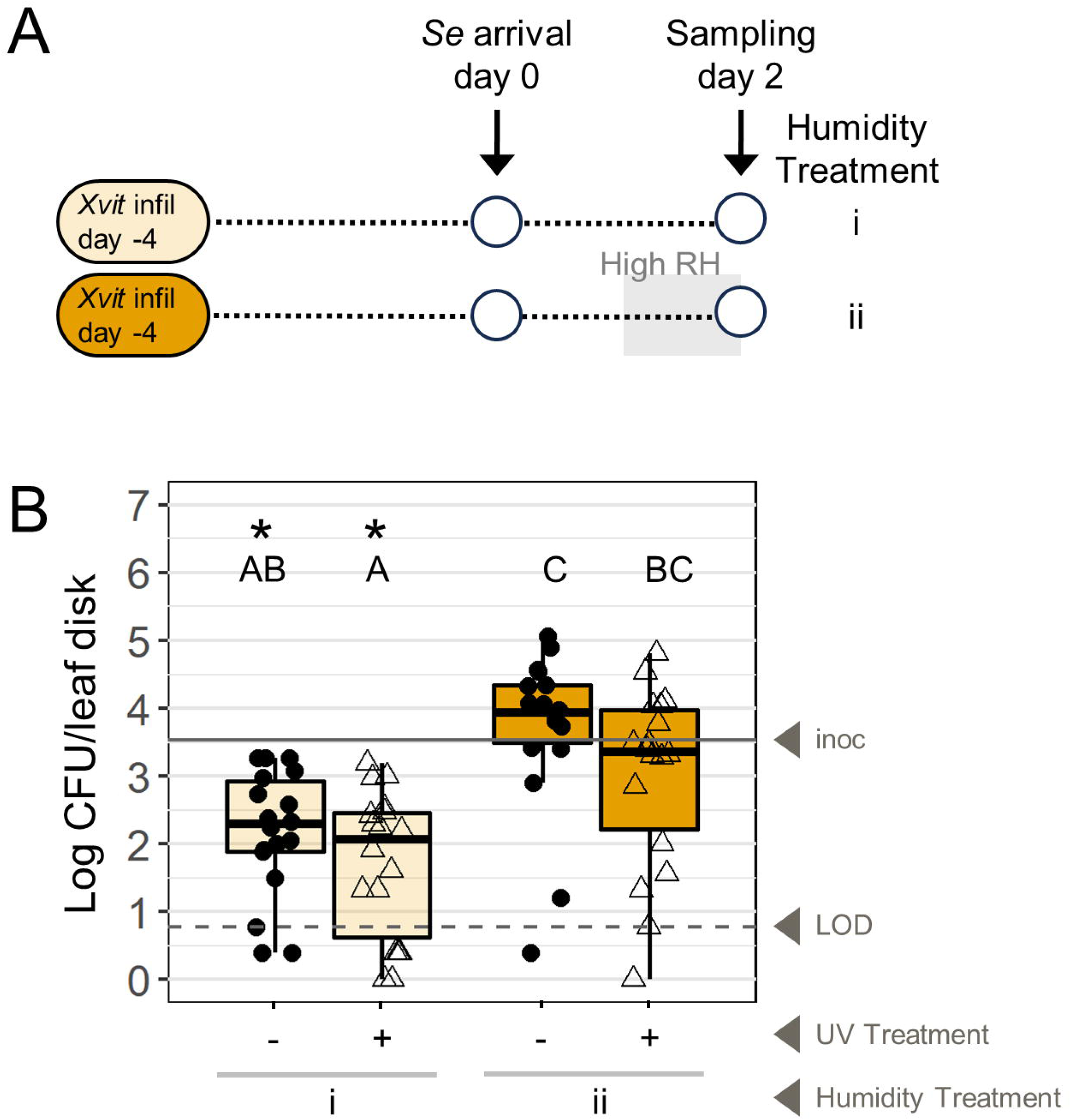
High humidity (>90% RH) exposure is not required for *S. enterica* to colonize UV-protected niches of *X. vitians*-infected lettuce. **(A)** Schematic of experimental design. Lettuce plants were divided into two treatment groups. *S. enterica* (*Se*) cells were applied to leaves previously infiltrated with *X. vitians* (*Xvit*) at 3 days post-infiltration. Plants were sampled following 0 or 24 hours of exposure to high humidity (>90% RH) prior to sampling (Humidity Treatments i and ii, respectively). **(B)** *S. enterica* colonization of *X. vitians*-infiltrated plants at 2 days post-arrival (dpa) in response to varying humidity treatments. Leaf halves were irradiated with UV immediately prior to sampling (unfilled triangle points) or left non-irradiated (filled circle points). Letters above the boxplots indicate significance between treatment groups based on Welch-ANOVA and *post hoc* Games-Howell means comparison tests (*P* < 0.05). Asterisks (*) indicate that *S. enterica in planta* colonization is significantly less than the arrival population based on one-sample Wilcoxon tests (*P* < 0.05). Solid line indicates average *S. enterica* arrival population applied to each sampled area, and dashed line indicates limit of detection (LOD). *N* = 18 leaves/treatment group; data is pooled from three independent experimental replicates.

Similarly to the previous experiment, high humidity (Humidity Treatment ii) resulted in higher *S. enterica* populations at 2 days post-*S. enterica* arrival on *X. vitians-*infected leaves (**Figure 3B**; *P =* 3.55 x 10^-5^, Welch ANOVA). *S. enterica* populations on plants that were not exposed to high humidity (Humidity Treatment i) were below the arrival population level at time of sampling. Although humidity had a significant impact on *S. enterica* persistence in general, for the viable *S. enterica* cells that were present, humidity did not significantly contribute to *S. enterica’s* ability to colonize UV-protected niches. Contrary to our hypothesis, UV irradiation did not cause a significant reduction in *S. enterica* populations for either humidity treatment, indicating that high humidity is not required for *S. enterica* colonization of UV-protected niches (**Figure 3B**). Notably, the introduction of high humidity at 5 dpi was found to have no effect on *X. vitians* populations and host electrolyte leakage in this experiment **(Supplementary** Figures 3A-C), demonstrating that the observed impact of humidity on *S. enterica* persistence is attributed to the humidity environment, not plant disease or *X. vitians*.

### Disease progression by *X. vitians* influences *S. enterica* ingress and survival within infected lettuce

We observed that *X. vitians* infection alters the host over time, with visible watersoaking onset occurring at 2 dpi (**Supplementary** Figure 5). During late infection (approximately 5 dpi), *X. vitians*-infected tissue began to appear drier, signaling the onset of tissue necrosis (**Supplementary** Figure 5). These observations prompted the question of whether an ideal window during *X. vitians* infection exists that optimizes *S. enterica* growth and apoplast colonization, which we addressed by varying the timing of *S. enterica’s* arrival on *X. vitians-*infected leaves relative to *X. vitians* infection progress (**Figure 4A**). *S. enterica* arrival on healthy leaves or on *X. vitians-*infected leaves at 1 dpi (Treatments H and X1) generally resulted in the lowest *S. enterica* populations after two days of *S. enterica* colonization (**Figure 4B**; non-irradiated leaves: *P* = 0.001, Welch ANOVA; UV-irradiated leaves: *P* = 1.61 x 10^-9^, Welch ANOVA). *S. enterica* populations recovered from both non-irradiated and UV-irradiated leaves for Treatments H and X1 were significantly lower than the *S. enterica* arrival population level (**Figure 4B**; *P* < 0.001, Wilcoxon signed rank test). *S. enterica* colonization of non-irradiated lettuce samples revealed that arrival of *S. enterica* at 1 dpi (Treatment X1) resulted in lower *S. enterica* populations on non-irradiated leaves than those on healthy (Treatment H) lettuce leaves, demonstrating that *S. enterica* arrival on *X. vitians-*infected leaves can result in greater population decline than that of healthy leaves if *S. enterica* arrival precludes the onset of *X. vitians-*induced watersoaking (**Figure 4B**). The absence of an aqueous apoplast in Treatments H and X1 also corresponded with the lowest *S. enterica* colonization of UV-protected niches at 2 days post-*S. enterica* arrival (**Figure 4B**).

**Figure 4:**
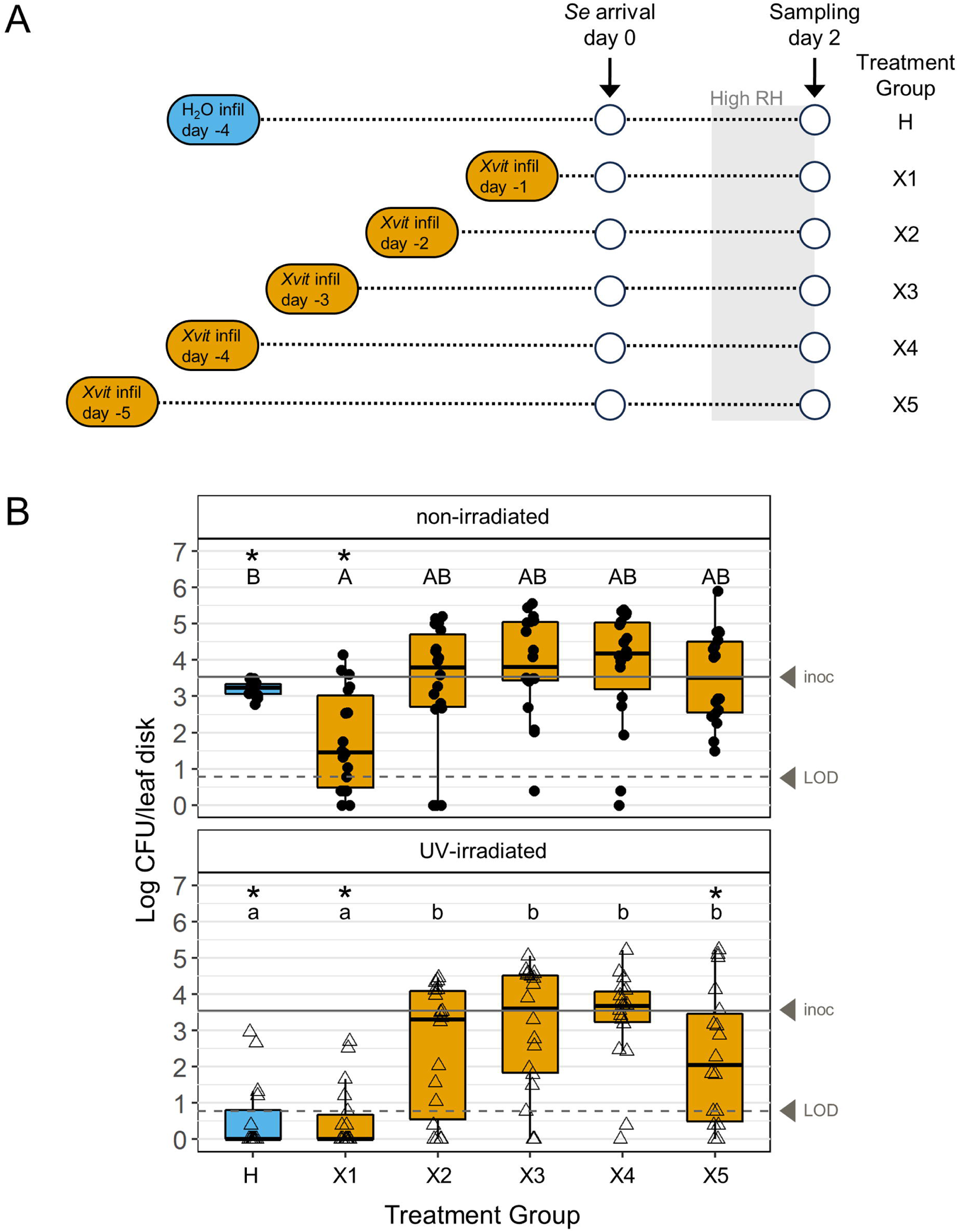
The fate of *S. enterica* on lettuce leaves is influenced by *X. vitians* infection progress at *S. enterica* arrival. **(A)** Schematic of experimental design. Lettuce plants were divided into six treatment groups. *S. enterica* (*Se*) cells were applied to leaves previously infiltrated with H_2_O or *X. vitians* (*Xvit*) at various days post-infiltration (dpi). Plants were sampled following 24 hours of exposure to high humidity (>90% RH). **(B)** *Se* colonization of H_2_O-infiltrated (blue boxplots, Treatment H) and *X. vitians*-infiltrated (orange boxplots, Treatments X#) plants at 2 days *post-Se* arrival (dpa). Leaf halves were irradiated with UV immediately prior to sampling (unfilled triangle points) or left non-irradiated (filled circle points). Letters above the boxplots indicate significance between treatment groups (H-X5) for non-irradiated leaves (uppercase) or UV-treated leaves (lowercase) based on Welch-ANOVA and *post hoc* Games-Howell means comparison tests (*P* < 0.05). Asterisks (*) indicate that *Se in planta* colonization is significantly less than the arrival population based on one-sample Wilcoxon tests (*P* < 0.05). Solid line indicates average *Se* arrival population applied to each sampled area, and dashed line indicates limit of detection (LOD). *N* = 15-18 leaves/treatment group; data is pooled from three independent experimental replicates.

UV irradiation had a significant impact on reducing *S. enterica* populations for Treatments H, X1, and X5, specifically (*P* = 4.76 x 10^-8^, *P* = 3.65 x 10^-3^, and *P* = 0.0216, respectively; Welch T-test) making evident that *S. enterica* colonization of the apoplast was limited and the majority of the population remained on the leaf surface vulnerable to UV irradiation. By contrast, UV irradiation treatment did not have a significant effect on the *S. enterica* populations for Treatments X2-X4 (*P* > 0.05, Welch T-test), demonstrating that *S. enterica* arrival prior to necrosis onset when the water-soaked infection zone appears to be most translucent (**Supplementary** Figure 5) promotes *S. enterica* colonization of UV-protected niches of a *X. vitians-*infected host. When *S. enterica* arrived between 2-4 dpi (Treatments X2-X4), *S. enterica* populations persisted at levels similar to the arrival population level on UV-irradiated leaves (**Figure 4B**), whereas arrival at 5 dpi resulted in UV-protected *S. enterica* populations that were lower than the arrival population (**Figure 4B**; *P* < 0.01, Wilcoxon signed rank test) suggesting arrival late in plant disease progress when tissue is necrotic limits *S. enterica* to the leaf surface.

Infection progress coincided with *S. enterica’s* ability to grow. We found that *X. vitians* populations and host electrolyte leakage had reached a maximum level during late infection (5-7 dpi, **Supplementary** Figures 6A-B). Arrival of *S. enterica* between 2-5 dpi resulted in the highest proportions of lettuce samples for which *S. enterica* colonization exceeded the arrival population, indicating high incidence rates of *S. enterica* replication (**Table 2**). For Treatment X4, arrival of *S. enterica* at 4 dpi resulted in 72% of non-irradiated samples and 61% of UV-irradiated samples being colonized with higher *S. enterica* populations than the arrival population (**Table 2**). Altogether, these results provide evidence that there is a window of time during *X. vitians* infection when *S. enterica* persistence and potential to colonize UV-protected niches is optimized—this window opens at water soaking onset and closes during necrosis onset later in disease progression.

**TABLE 2:** Incidence*^a^* of *S. enterica in planta* growth depending on *X. vitians* infection progress at time of *S. enterica* arrival.

| UV Treatment Group<sup>c</sup> | Infiltration Treatment Group<sup>b</sup> |  |  |  |  |  |
| --- | --- | --- | --- | --- | --- | --- |
|  | H | X1 | X2 | X3 | X4 | X5 |
| Non-irradiated | 7% A<sup>d</sup> | 17% AB | 56% BC | 56% BC | 72% C | 50% BC |
| UV-irradiated | 0% a | 0% a | 33% ab | 50% b | 61% b | 28% ab |

<sup>a</sup>Percentage of samples for which in *planta* *S. enterica* colonization exceeds the arrival population.
<sup>b,c</sup>Treatment groups refer directly to experiment described in **Figure 2**.
<sup>d</sup>Letters indicate significance between treatment groups among non-irradiated leaves (uppercase) and among UV-treated leaves (lowercase) based on Fisher's exact tests ( $P < 0.05$ ).

Confocal laser scanning microscopy (CLSM) was utilized to visualize *S. enterica* localization *in planta* on the lettuce surface (**Figure 5A-F**) and within the apoplast (**Figure 5G-L**). For H_2_O-infiltrated leaves at 4 dpi, *S. enterica* cells were found only on the surface. *S. enterica* cells were observed colonizing at least one stomatal opening (z-depth = 0) for 56% (*N* = 9) of images fields obtained from H_2_O-infiltrated leaves (**Figure 5D**, for example), yet *S. enterica* failed to colonize the apoplast (**Figure 5J**, for example) in all these cases. By contrast, *S. enterica* colonization within substomatal areas of the apoplast was observed for *X. vitians*-infected leaves. Congruous to *in planta S. enterica* colonization results that showed *S. enterica* colonization of the UV-protected niches of *X. vitians*-infected leaves was higher following arrival at 4 dpi compared to 1 dpi (**Figure 4B**), CLSM revealed greater *S. enterica* colonization of the apoplast during late infection compared to early infection (**Figure 5H**, early infection; **Figure 5I**, late infection). *S. enterica* arrival at 4 dpi resulted in relatively frequent colonization of substomatal spaces within the apoplast, with 73% of image fields showing evidence of *S. enterica* ingress and only 27% of image fields showing an absence of *S. enterica* ingress (*N* = 15). (**Figure 5L**). For *S. enterica* arrival at 4 dpi, 87% of image fields (*N* = 15) included *S. enterica* colonization of at least one stomatal opening. In comparison, arrival of *S. enterica* on *X. vitians-*infected leaves as early at 1 dpi resulted in rare apoplast colonization (15% of fields imaged; *N* = 13; **Figure 5K**), despite 100% of fields showing *S. enterica* cells colonizing at least one stomatal opening (*N* = 13). **Figure 5E** representatively illustrates this, with three stomatal openings colonized by *S. enterica* cells at the surface, but only one of those same three stomata containing *S. enterica* within its substomatal space (**Figure 5K**). Our *S. enterica* population data (**Figure 4B**) and CLSM results altogether support that *S. enterica* arrival to leaves with established *X. vitians* infection promotes *S. enterica* colonization of the UV-protected apoplast.

**Figure 5:**
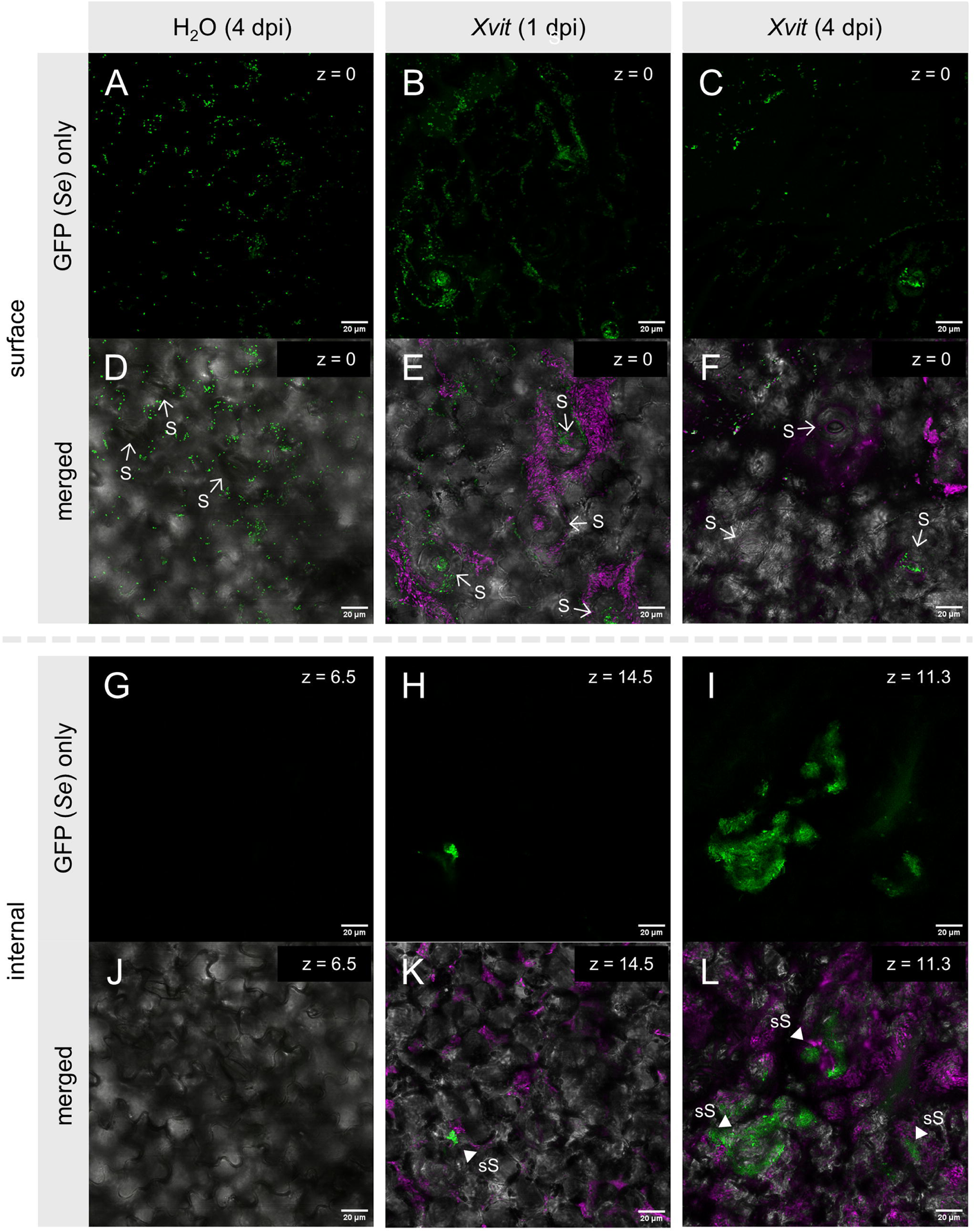
*S. enterica* colonizes the apoplast of *X.* vitians-infected lettuce leaves. CLSM images depict GFP-labelled *S. enterica* colonization of the lettuce surface. **(A-F)** and apoplast (**G-L**) for three sites **(columns A-C)** representative of three treatments and imaged at the given z-depths (μm). *S. enterica* cells were applied to the surface of leaves previously infiltrated with H_2_O (column A) or mCherry-labelled *X. vitians* cells **(columns B-C)**. Horizontal labels describe infiltration treatment and days-post infiltration (dpi) at time of *S. enterica* arrival. **A-C, G-I**: GFP channel only. **D-F, J-L**: merged channels (GFP, green; mCherry, magenta; DIC, bright-field). Stomata (S) and substomatal *S. enterica* cells (sS) are highlighted with white arrows and white triangles. Scale bar = 20 μm.

### *S. enterica* grows in a sucrose-rich environment in the presence of *X. vitians*

Sucrose is one of the most prevalent carbohydrate sources available in plant tissues (36), yet sucrose is a poor carbon source for *S. enterica* and not utilizable by most *S. enterica* strains (37). Having observed that *S. enterica* successfully replicates *in planta* in the presence of an established *X. vitians* infection (**Figures 1-2, Table 2**), we hypothesized that *X. vitians* catabolism of plant compounds directly aids in *S. enterica* replication via 1) *X. vitians*-associated compounds or 2) compounds secreted by *X. vitians.* To test this hypothesis, we developed an *in vitro* experiment in which *S. enterica* cells were introduced to turbid *X. vitians* cultures that had grown in M9 minimal media containing only sucrose as a carbon source (M9 sucrose) for 24 or 48 hours. Cell-free supernatant (CFS) was produced from the *X. vitians* cultures and introduced to *S. enterica* to compare the effect of intact *X. vitians* cells versus media containing secreted compounds. In addition, *S. enterica* cells were added to M9 sucrose or M9 glucose media alone as negative and positive controls.

We found that *S. enterica* cells incubated in co-culture with *X. vitians* cells or CFS replicated to population levels that exceeded the initial cell concentrations after 28 hours of incubation (**Figure 6**; *P =* 1.01 x 10^-7^, Welch ANOVA). By contrast, *S. enterica* incubation in M9 sucrose in monoculture (negative control; light blue bar) resulted in no growth after 28 hours. Specifically, introduction of *S. enterica* to *X. vitians* cells that had been growing in M9 sucrose for 24 hours (yellow bar) resulted in *S. enterica* cell concentrations that were greater than the negative control and greater than either CFS treatment (emerald and dark green bars), showing evidence for bacterial cross-feeding supporting *S. enterica* growth. Meanwhile, introducing *S. enterica* to a 48-hour *X. vitians* culture (orange bar) resulted in higher variability in *S. enterica* cell concentration than the 24-hour culture (yellow bar), and resulted in similar *S. enterica* concentrations to both the negative control and CFS treatments. Notably, the 24-hour *X. vitians* culture exhibited additional *X. vitians* growth during the 28-hour period (**Supplementary** Figure 7), whereas the 48-hour culture did not, suggesting that replicating *X. vitians* cells confer the greatest nutritional benefit to *S. enterica*.

**Figure 6:**
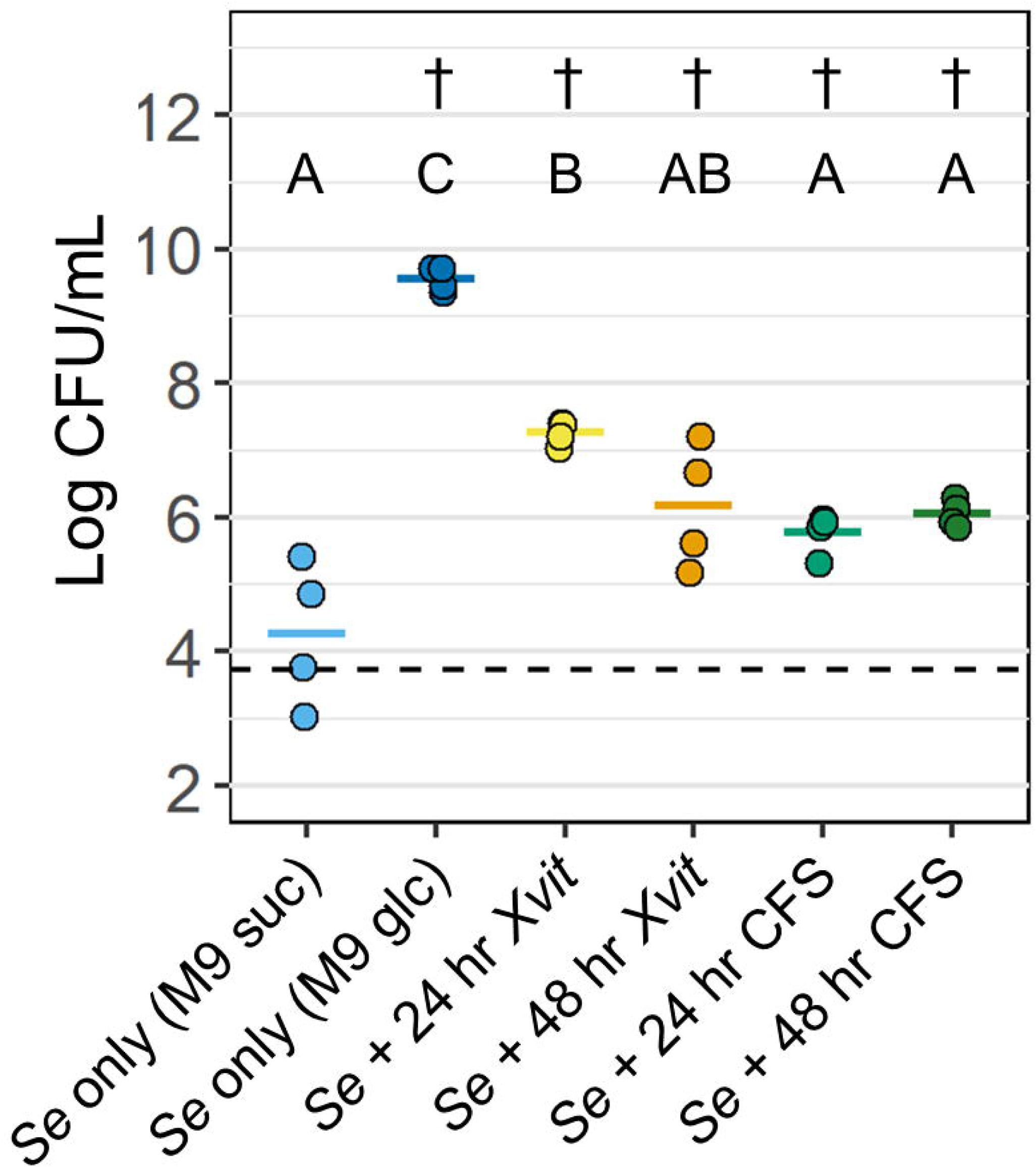
*X. vitians* metabolism of sucrose produces nutrients *S. enterica* can utilize for growth. *S. enterica* cells were added to fresh M9 sucrose (M9 suc, light blue points) or M9 glucose (M9 glc, dark blue points), mixed with *X. vitians* (*Xvit*) cell cultures grown for 24 (yellow points) or 48 hours (orange points) in M9 sucrose, or mixed with *Xvit* cell-free supernatant (CFS, emerald green & dark green points) at time (t) = 0 hr. Dotted line represents the initial *S. enterica* cell concentration at t = 0 hr. Each horizontal colored bar represents the average *S. enterica* cell concentrations of four independent experiments at t = 28 hr. Letters above the boxplots indicate significance between treatment groups based on Welch-ANOVA and *post hoc* Games-Howell means comparison tests (*P* < 0.05). Daggers (†) indicate a significant increase in *S. enterica* cell concentration compared to t = 0 hr based on two-sample T-tests (P < 0.05).

## DISCUSSION

Water—a habitat, vehicle for bacteria movement, and vital source of moisture needed for growth—influences *S. enterica’s* lifestyle in agricultural environments. Our study, which involved modelling the arrival of *S. enterica* on the surface of leaves via splash dispersal (**Figures 1-4**) and investigates the interactions between phytobacterial infection and humidity, largely relates to water. Irrigation water is a common reservoir for *S. enterica* (21, 38, 39) and has been previously implicated in enteric pathogen outbreaks associated with raw produce (40, 41). *S. enterica’s* presence in irrigation water (40, 42) offers the potential for *S. enterica* to migrate to leaves and fruit. Sprinkler and microsprinkler irrigation are commonly used for agricultural fields in the United States (44, 45), including California and Arizona where most leafy greens are cultivated (46). However, this form of irrigation has inherent risk. Splash dispersal has been implicated as a vehicle for microbial dispersal on plant foliage (47, 48). In this study, we investigated the relationship (49, 50) between an enteric pathogen and a plant pathogen colonizing a lettuce host and examined how humidity may act on the enteric pathogen’s ability to benefit from plant disease. *S. enterica* was found to thrive on lettuce plants affected by bacterial spot disease (**Figures 1B-C, 2B,** and **4B**). The potential for *S. enterica* to survive and replicate on lettuce plants was linked to *X. vitians* infection progress (**Figure 4B**) and exposure to high humidity **(Figure 2B**), with our results showing that the combination of water soaking caused by disease and high humidity supported the highest *S. enterica* populations *in planta.* Lastly, by measuring *S. enterica* growth in a sucrose-based minimal media, we identified bacterial cross-feeding as a nutrient source for *S. enterica* in association with *X. vitians* **(Figure 6**).

Humidity is recognized as a driver of plant-bacterial interactions (51). High humidity has been shown to correlate with increases in phytobacterial disease (3, 52), including bacterial spot caused by *X. vitians* (53), and linked to the compromise of stomatal-based defense (54). In addition, enteric pathogen populations such as *S. enterica* have been shown to exhibit slower population decline on healthy plants exposed to high humidity (55–58). To determine how humidity impacts *S. enterica’s* ability to benefit from *X. vitians* disease, we tested the impact of a 24-hour and 72-hour exposure to >90% RH conditions (**Figure 2A**). Our data showed that *S. enterica* populations were higher on *X. vitians-*infected plants exposed to high humidity, yet unaffected by humidity on healthy plants **(Figure 2B**). The latter result that high humidity did not have a significant impact on *S. enterica* persistence on healthy plants is somewhat contrary to published literature demonstrating that high humidity supports *S. enterica* survival on healthy plants (55–58). However, the *S. enterica* inoculum used for this study, ∼3000 CFU/droplet, is relatively low compared to these previous studies examining humidity impacts on *S. enterica,* and lower inoculum can result in greater variability (57). Furthermore, it is worth emphasizing that the effect of high humidity on enhancing *S. enterica* growth in the presence of disease was not linked directly to *X. vitians* cell density. Unexpectedly, we found that the introduction of high humidity to *X. vitians*-infected leaves could inhibit *X. vitians* colonization (**Supplementary** Figure 2C), though this effect was not observed when the high humidity conditions were introduced later during infection (**Supplementary** Figure 3C). This observation about humidity and *X. vitians* infection is contrary to the well-known association between high humidity and phytopathogenic disease, including disease by *X. vitians* (3, 52). However, these particular experiments only looked at the short-term impact of high humidity on *X. vitians* infection. Short-term disruptions to *X. vitians* infection progress may not necessarily predict how high humidity would impact infection of the course of multiple days or even weeks. Though *S. enterica* and *X. vitians* exhibited differing responses to a 24-hour high humidity exposure, the long-term impact of high humidity climate trends on *S. enterica* colonization of plants in association with phytobacterial disease is unknown. Considering that precipitation and seasonality have been shown to influence *S. enterica’s* abundance in bodies of water, including irrigation ponds (43, 59, 60), the question of how humidity impacts *S. enterica* in the field is worth exploring. Plant disease is also influenced by humidity and season, and it is plausible that *S. enterica* prevalence on plants is influenced by humidity patterns as well.

Our previous study, which also modelled a contamination route via splash dispersal, found that *S. enterica* applied to tomato leaves via surface droplet could become internalized into the leaf apoplast (17). A wet apoplast, either abiotically water-logged or water-soaked from phytobacterial disease, could enable the physical absorption of surface water and passive ingress of *S. enterica* in a non-flagellar motility-dependent manner (17). This study expands on those findings by examining the role of relative humidity in *S. enterica’s* ability to persist on infected leaves and colonize the apoplast. In the absence of high humidity exposure that facilitates replication, *S. enterica* still managed to colonize the UV-protected apoplast of *X. vitians-*infected lettuce (**Figure 3B**), demonstrating that *S. enterica’s* ability to replicate under high humidity conditions is not explained by colonization of the apoplast alone. Although moderate humidity conditions (50-70%) were not conducive to *S. enterica* replication in the presence of *X. vitians* infection (**Figure 3B**), *X. vitians* replicated successfully *in planta* at 50-70% RH and caused measurable increases in electrolyte leakage (**Supplementary** Figures 4A-B) suggesting that moderate humidity conditions can allow the liberation of apoplast nutrients during *X. vitians* infection. Therefore, these results support the idea that the combination of extended high humidity and a *X. vitians-*infected apoplast provides additional resources *S. enterica* can successfully metabolize. Whether high humidity conditions must be sustained continuously or if the accumulation of multiple high humidity events is sufficient to promote *S. enterica* growth on *Xanthomonas-*infected leaves should be determined through future work.

Stomata serve as critical entry points for bacteria in the phyllosphere (61). *Xanthomonas* infection has been previously shown to promote ingress of surface bacteria into the apoplast of tomato leaves via stomata (17). In this study, we observed that *S. enterica* became internalized within *X. vitians-*infected lettuce leaves, based on evidence of *S. enterica* protection from UV irradiation (**Figures 1, 3-4**) and colonization of *S. enterica* within substomatal chambers of the apoplast (**Figure 5**). If a bacterium can enter the apoplast via stomata and cells are found on the leaf surface at stomata, one might assume that they would also be found in the substomatal cavity. We found with CLSM that *S. enterica* ingress varied among stomata. *S. enterica* colonization of stomatal openings, likely influenced by *S. enterica* chemotaxis (62), did not always result in colonization of the lower substomatal space (**Figure 5**). Syringe infiltration of *X. vitians* into the apoplast is assumed to create a consistent dispersal of cells within the infiltration zone (63). However, the mottled appearance of the infected area suggests that disease onset is not consistent throughout the inoculation zone (**Supplementary** Figure 5). Clonal *Pseudomonas syringae* cells have been shown to exhibit heterogeneity in T3SS expression within the apoplast (64). Our CLSM results showed a heterogeneity of *X. vitians* microcolonies on the leaf surface and within the apoplast, and we speculate that this heterogeneity may differentially impact *S. enterica’s* interactions with stomata and colonization of the apoplast.

*S. enterica* was also influenced by the temporal dynamics of *X. vitians* disease progress. Our study investigated the fate of *S. enterica* following arrival at various times during *X. vitians* infection progress and essentially used *S. enterica* as a biological reporter for changes to a *X. vitians-*infected apoplast. Although *X. vitians* infection was found to be generally beneficial to *S. enterica*, (**Figures 1-2, 5**), the early infection court was a hostile environment for *S. enterica* (**Figure 4**). Arrival of *S. enterica* during early infection at 1 dpi led to a reduction in *S. enterica* populations, sometimes to non-detectable levels two days later (**Figure 4**, Treatment X1). We hypothesize that *S. enterica’s* population reduction and death in the host experiencing an early *X. vitians* infection is attributed to the host defense response. Infiltration of bacteria into the apoplast can elicit MAMP (microbe-associated molecular pattern)-triggered immune (MTI) response (65). Recognition of phytobacterial invaders may trigger phytohormone signaling and apoplast accumulation of reactive oxygen and nitrogen species (66, 67), which has been demonstrated to limit *S. enterica* growth *in planta* (68). *S. enterica* flagellin is a known elicitor of MTI (69), and *S. enterica* may benefit from suppression of MTI by phytobacteria based on its ability to thrive in the presence of disease (17). Hence, modulation of plant defenses may contribute to *S. enterica* success on infected leaves. However, as demonstrated in our study, the timing of co-colonization is important. Previously, it was shown that *S. enterica* achieved greater colonization of tomato foliage when seeds were co-treated with *Xanthomonas euvesicatoria* (formerly known as *X. campestris* pv. vesicatoria) via inoculated soil (18). However, at the early tomato seedling stage, *S. enterica* populations were lower in the presence of *X. euvesicatoria*. Additionally, Potnis et al. (70) demonstrated that *S. enterica* persistence on tomato plants is enhanced by *Xanthomonas perforans* infection but diminished by an avirulent *X. perforans* strain that can neither cause disease, nor suppress MTI. Hence, *S. enterica* responds to both aggravation and suppression of host defenses by phytobacteria.

Phytobacterial infection likely also supports *S. enterica* survival in the phyllosphere by providing nutrients for growth. *S. enterica* metabolism is incredibly flexible, enabling it to utilize many different types of carbon and nitrogen sources in an animal host, even at scant concentrations (71). However, *S. enterica* growth is disadvantaged on plants. *S. enterica* cannot liberate nutrients from plant cells on its own because it lacks cell-wall degrading enzymes and the ability to manipulate the host environment via type-III effectors (72–74). Additionally, as previously mentioned, *S. enterica* is rarely able to utilize sucrose, one of the most abundant carbon sources on plants (75, 76). *S. enterica* can persist in the phyllosphere for months on healthy plants (22), though its population levels steadily decline (16, 22, 77), presumably due to stress exposure. Contrastingly, in the presence of *X. vitians* infection, *S. enterica* replicates *in planta* (**Figures 1-2**, **Table 2**), indicating that infection increases nutrient availability for *S. enterica*. However, as *X. vitians* infection leads to tissue necrosis (**Supplementary** Figure 2) and presumably a decrease in available nutrients, *S. enterica* populations in a *X. vitians-*infected apoplast decline, as evident by *S. enterica* populations dropping below the original arrival population level following arrival during at 5 dpi (**Figure 4B**, Treatment X5). Late in infection, phytobacterial pathogens migrate away from necrotic tissue, presumably to locate available nutrients or to disperse to another host. Future studies will examine the dispersal of *S. enterica* beyond initially infected leaf tissue to determine if the human and plant pathogens co-migrate as they seek nutrient sources.

Nutrients metabolized by *S. enterica* in the presence of disease are likely host derived. *Xanthomonas* replication in the apoplast during infection is driven by leakage of plant cell constituents (8, 16), which *S. enterica* may be able to scavenge in the presence of *X. vitians* infection. However, we also hypothesize that bacterial cross-feeding (78) between *S. enterica* and *X. vitians* contributes to *S. enterica’s* ability to grow *in planta.* Previously, cross-feeding has been demonstrated to influence nutritional niches, thus affecting phytobacterial community composition (79). Our finding that infection can promote *S. enterica* replication within the UV-protected apoplast (**Figure 1C**) and our CLSM observations of *S. enterica* cells localized in proximity to *X. vitians* cells *in planta* (**Figure 5**) inspired us to test whether these cells interact directly via nutrient flow from *X. vitians* to *S. enterica.* The results of our *in vitro* experiment demonstrated that although sucrose is a poor nutrient source for *S. enterica*, *S. enterica* could grow in sucrose-based minimal media if replicating *X. vitians* cells were present (**Figure 6**).

Cell-free supernatant (CFS) derived from *X. vitians* cultures did not support *S. enterica* growth as well as cultures containing *X. vitians* cells. *Xanthomonas* strains are known for producing sticky extracellular polysaccharides, including xanthan gum (80), which may have contributed to the exclusion of nutrients during the filtering of *X. vitians* culture supernatant. The identity of the nutrients utilized by *S. enterica* via cross-feeding are unknown, but could include amino acids, carbohydrates, or organic acids (75). Future studies may elucidate the carbon sources metabolized by *S. enterica in planta* in the presence of disease.

Previous work investigating relationships between enteric pathogen persistence, phytopathogen infection, and environment has generally studied 1-2 of these elements in isolation. In reality, interplay between enteric pathogens, plant pathogens, and climate factors are all likely to influence the microbial communities in the phyllosphere. In this study expanding knowledge of *S. enterica’s* fate following water droplet dispersal, we found that *S. enterica* survival on lettuce is enhanced by bacterial spot disease caused by *X. vitians* and that *S. enterica* proliferation was conditional on the status of infection and the environment **(Figure 7**). Overall, it was the combination of high humidity exposure and the timing of *S. enterica* arrival with respect to bacterial spot disease progress that maximized *S. enterica* survival and access to nutrients in an otherwise inhospitable host. As climate patterns continue to evolve, microbial dynamics in the phyllosphere will evolve too, underlining the importance of studying host-microbial interactions under various environmental conditions.

**Figure 7:**
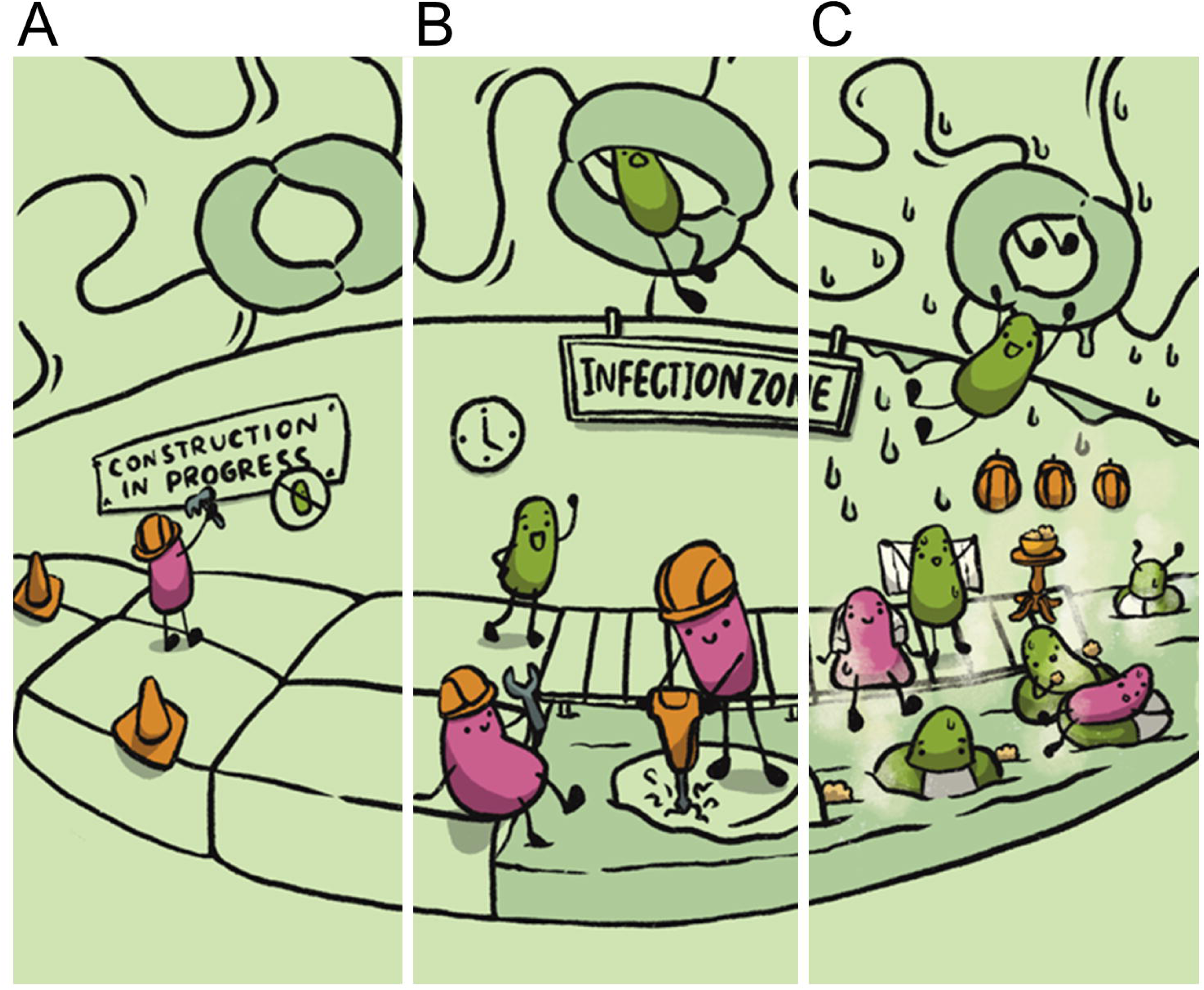
The fate of *Salmonella enterica* in the phyllosphere is influenced by *Xanthomonas* infection progress and humidity. **(A)** Early *Xanthomonas* infection. *X. vitians* (magenta bacteria) begin to modify the host environment, but the infection court is not yet habitable for *S. enterica.* **(B)** Established *Xanthomonas* infection. *S. enterica* (green bacteria) successfully colonize the apoplast of *Xanthomonas-*infected leaves via ingress through stomata. **(C)** *Xanthomonas* infection plus high humidity. High humidity (>90% RH) promotes replication of *S. enterica* within a *Xanthomonas-*infected apoplast, though is not required for *S. enterica* ingress.

## ACKNOWLEDGMENTS

This material is based upon work supported by the National Science Foundation Graduate Research Fellowship under grant no. DGE-2137424 and by the UW-Madison Food Research Institute. Any opinions, findings, and conclusions or recommendations expressed in this material are those of the authors and do not necessarily reflect the views of the National Science Foundation. The funders had no role in study design, data collection, and interpretation or the decision to submit the work for publication.

Thank you to J. Jones’ and Z. Gitai’s labs for sharing bacterial strains that helped make this work possible. We especially thank N. Ouzounov for constructing the pNO-mCherry plasmid used in this study. We thank C. Solis-Lemus for statistical consultation of the entire study and mentoring M.H.D. in statistical analysis.

Confocal microscopy was performed at the Newcomb Imaging Center, Department of Botany, University of Wisconsin, Madison, WI. We thank S. Swanson for her training and assistance with CLSM.

We would also like to acknowledge H. Pai, whose illustration work featured in (81) provided inspiration for our cartoon summary figure.

M.H.D and J.D.B. developed the experiments. M.H.D., D.N., and S.C.Z. performed the experiments. M.H.D. and J.D.B. wrote the manuscript, and all authors reviewed the manuscript. M.H.D. created the original illustrations.

## REFERENCES

1. Schlechter RO, Miebach M, Remus-Emsermann MNP. 2019. Driving factors of epiphytic bacterial communities: A review. J Adv Res 19:57–65.

2. Vorholt JA. 2012. Microbial life in the phyllosphere. Nat Rev Microbiol 10:828–840.

3. Xin XF, Nomura K, Aung K, Velásquez AC, Yao J, Boutrot F, Chang JH, Zipfel C, He SY. 2016. Bacteria establish an aqueous living space in plants crucial for virulence. Nature 539:524–529.

4. Hu Y, Ding Y, Cai B, Qin X, Wu J, Yuan M, Wan S, Zhao Y, Xin X-F. 2022. Bacterial effectors manipulate plant abscisic acid signaling for creation of an aqueous apoplast. Cell Host Microbe 1–12.

5. Roussin-Léveillée C, Lajeunesse G, St-Amand M, Veerapen VP, Silva-Martins G, Nomura K, Brassard S, Bolaji A, He SY, Moffett P. 2022. Evolutionarily conserved bacterial effectors hijack abscisic acid signaling to induce an aqueous environment in the apoplast. Cell Host Microbe 1–13.

6. Schornack S, Minsavage G V, Stall RE, Jones JB, Lahaye T, Jones JB. 2006. Characterization of AvrHah1, a novel AvrBs3-like effector from Xanthomonas gardneri with virulence and avirulence activity. New Phytologist 179:546–556.

7. Cowles KN, Block AK, Barak JD. 2022. Xanthomonas hortorum pv. gardneri TAL effector AvrHah1 is necessary and sufficient for increased persistence of Salmonella enterica on tomato leaves. Sci Rep 12:1–13.

8. Schwartz AR, Morbitzer R, Lahaye T, Staskawicz BJ. 2017. TALE-induced bHLH transcription factors that activate a pectate lyase contribute to water soaking in bacterial spot of tomato. Proceedings of the National Academy of Sciences 114:E897–E903.

9. Dia NC, Morinière L, Cottyn B, Bernal E, Jacobs JM, Koebnik R, Osdaghi E, Potnis N, Pothier JF. 2022. *Xanthomonas hortorum* – beyond gardens: Current taxonomy, genomics, and virulence repertoires. Mol Plant Pathol 23:597–621.

10. Robinson PE, Jones JB, Pernezny K. 2006. Bacterial leaf spot of lettuce: Relationship of temperature to infection and potential host range of Xanthomonas campestris pv. vitians. Plant Dis 90:465–470.

11. Savary S, Willocquet L, Pethybridge SJ, Esker P, McRoberts N, Nelson A. 2019. The global burden of pathogens and pests on major food crops. Nat Ecol Evol 3:430–439.

12. Goudeau DM, Parker CT, Zhou Y, Sela S, Kroupitski Y, Brandl MT. 2013. The Salmonella transcriptome in lettuce and cilantro soft rot reveals a niche overlap with the animal host intestine. Appl Environ Microbiol 79:250–262.

13. Kwan G, Charkowski AO, Barak JD. 2013. Salmonella enterica Suppresses Pectobacterium carotovorum subsp. carotovorum Population and Soft Rot Progression by Acidifying the Microaerophilic Environment. mBio 4:1–9.

14. Meng F, Altier C, Martin GB. 2013. Salmonella colonization activates the plant immune system and benefits from association with plant pathogenic bacteria. Environ Microbiol 15:2418–2430.

15. Potnis N, Soto-Arias JP, Cowles KN, van Bruggen AHC, Jones JB, Barak JD. 2014. Xanthomonas perforans Colonization Influences Salmonella enterica in the Tomato Phyllosphere. Appl Environ Microbiol 80:3173–3180.

16. Potnis N, Colee J, Jones JB, Barak JD. 2015. Plant Pathogen-Induced Water-Soaking Promotes Salmonella enterica Growth on Tomato Leaves. Appl Environ Microbiol 81:8126–8134.

17. Dixon MH, Cowles KN, Zaacks SC, Marciniak IN. 2022. Xanthomonas Infection Transforms the Apoplast into an Accessible and Habitable Niche for Salmonella enterica. Appl Environ Microbiol.

18. Barak JD, Liang AS. 2008. Role of soil, crop debris, and a plant pathogen in Salmonella enterica contamination of tomato plants. PLoS One 3.

19. Teplitski M, Moraes M De. 2018. Of Mice and Men and Plants: Comparative Genomics of the Dual Lifestyles of Enteric Pathogens. Trends Microbiol 26:748–754.

20. Spector MP, Kenyon WJ. 2012. Resistance and survival strategies of Salmonella enterica to environmental stresses. Food Research International 45:455–481.

21. Liu H, Whitehouse CA, Li B. 2018. Presence and Persistence of Salmonella in Water: The Impact on Microbial Quality of Water and Food Safety. Front Public Health 6.

22. Islam M, Morgan J, Doyle MP, Phatak SC, Millner P, Jiang X. 2004. Persistence of *Salmonella enterica* Serovar Typhimurium on Lettuce and Parsley and in Soils on Which They Were Grown in Fields Treated with Contaminated Manure Composts or Irrigation Water. Foodborne Pathog Dis 1:27–35.

23. Islam M, Morgan J, Doyle MP, Phatak SC, Millner P, Jiang X. 2004. Fate of Salmonella enterica Serovar Typhimurium on Carrots and Radishes Grown in Fields Treated with Contaminated Manure Composts or Irrigation Water. Appl Environ Microbiol 70:2497– 2502.

24. Brandl MT. 2006. Fitness of Human Enteric Pathogens on Plants and Implications for Food Safety. Annual Reviews of Phytopathology 44.

25. Carstens CK, Salazar JK, Darkoh C. 2019. Multistate Outbreaks of Foodborne Illness in the United States Associated With Fresh Produce From 2010 to 2017. Front Microbiol. Frontiers Media S.A. 10.3389/fmicb.2019.02667.

26. Horby PW, O’Brien SJ, Adak GK, Graham C, Hawker JI, Hunter P, Lane C, Lawson AJ, Mitchell RT, Reacher MH, Threlfall EJ, Ward LR. 2003. A national outbreak of multi-resistant *Salmonella enterica* serovar Typhimurium definitive phage type (DT) 104 associated with consumption of lettuce. Epidemiol Infect 130:169–178.

27. Lienemann T, Niskanen T, Guedes S, Siitonen A, Kuusi M, Rimhanen-Finne R. 2011. Iceberg Lettuce as Suggested Source of a Nationwide Outbreak Caused by Two Salmonella Serotypes, Newport and Reading, in Finland in 2008. J Food Prot 74:1035–1040.

28. McClure M, Whitney B, Gardenhire I, Crosby A, Wellman A, Patel K, McCormic ZD, Gieraltowski L, Gollarza L, Low MSF, Adams J, Pightling A, Bell RL, Nolte K, Tijerina M, Frost JT, Beix JA, Boegler KA, Dow J, Altman S, Wise ME, Bazaco MC, Viazis S. 2023. An Outbreak Investigation of Salmonella Typhimurium Illnesses in the United States Linked to Packaged Leafy Greens Produced at a Controlled Environment Agriculture Indoor Hydroponic Operation – 2021. J Food Prot 86:100079.

29. Ouzounov N. 2015. Establishment and maintenance of rod shape by the bacterial actin homologue MreB. Princeton University, Princeton, NJ.

30. Hashimoto-Gotoh T, Yamaguchi M, Yasojima K, Tsujimura A, Wakabayashi Y, Watanabe Y. 2000. A set of temperature sensitive-replication/-segregation and temperature resistant plasmid vectors with different copy numbers and in an isogenic background (chloramphenicol, kanamycin, lacZ, repA, par, polA). Gene 241:185–191.

31. Lutz R, Bujard H. 1997. Independent and tight regulation of transcriptional units in escherichia coli via the LacR/O, the TetR/O and AraC/I1-I2 regulatory elements. Nucleic Acids Res 25:1203–1210.

32. Dixon MH, Cowles KN, Zaacks SC, Marciniak IN. 2022. Xanthomonas Infection Transforms the Apoplast into an Accessible and Habitable Niche for Salmonella enterica. Appl Environ Microbiol.

33. Cowles KN, Groves RL, Barak JD. 2018. Leafhopper-Induced Activation of the Jasmonic Acid Response Benefits Salmonella enterica in a Flagellum-Dependent Manner. Front Microbiol 9:1–15.

34. Harrod VL, Groves RL, Guillemette EG, Barak JD. 2022. Salmonella enterica changes Macrosteles quadrilineatus feeding behaviors resulting in altered S. enterica distribution on leaves and increased populations. Sci Rep 12:1–13.

35. Kessens R, Sorensen N, Kabbage M. 2018. An inhibitor of apoptosis (SfIAP) interacts with SQUAMOSA promoter-binding protein (SBP) transcription factors that exhibit pro-cell death characteristics. Plant Direct 2:1–17.

36. Fatima U, Senthil-Kumar M. 2015. Plant and pathogen nutrient acquisition strategies. Front Plant Sci 6:1–12.

37. Reid SJ, Abratt VR. 2005. Sucrose utilisation in bacteria: Genetic organisation and regulation. Appl Microbiol Biotechnol 67:312–321.

38. Steele M, Odumeru J. 2004. Irrigation Water as Source of Foodborne Pathogens on Fruit and Vegetables. J Food Prot 67:2839–2849.

39. Jacobsen CS, Bech TB. 2012. Soil survival of Salmonella and transfer to freshwater and fresh produce. Food Research International 45:557–566.

40. Centers for Disease Control and Prevention (CDC). 2008. Outbreak of Salmonella Serotype Saintpaul Infections Associated with Multiple Raw Produce Items---United States, 2008. Morbidity and Mortality Weekly Report 57:929–934.

41. Centers for Disease Control and Prevention (CDC). 2018. Surveillance for Foodborne Disease Outbreaks United States, 2016, Annual Report. Atlanta, Georgia.

42. Centers for Disease Control and Prevention (CDC). 2018. Multistate Outbreak of E. coli O157:H7 Infections Linked to Romaine Lettuce (Final Update). https://www.cdc.gov/ecoli/2018/o157h7-04-18/index.html. Retrieved 11 January 2022.

43. Haley BJ, Cole DJ, Lipp EK. 2009. Distribution, Diversity, and Seasonality of Waterborne Salmonellae in a Rural Watershed. Appl Environ Microbiol 75:1248–1255.

44. Johnson R, Cody BA. 2015. California Agricultural Production and Irrigated Water Use.

45. Mpanga IK, Idowu OJ. 2021. A Decade of Irrigation Water use trends in Southwestern USA: The Role of Irrigation Technology, Best Management Practices, and Outreach Education Programs. Agric Water Manag 243:106438.

46. Davis W, Weber C, Wechsler S, Lucier G, Soria Rodriguez M, Yeh A, Fan X. 2023. Vegetable and Pulses Outlook: April 2023.

47. Hirano SS, Baker LS, Upper CD. 1996. Raindrop Momentum Triggers Growth of Leaf-Associated Populations of Pseudomonas syringae on Field-Grown Snap Bean Plants. Appl Environ Microbiol 62:2560–2566.

48. Kim S, Park H, Gruszewski HA, Schmale DG, Jung S. 2019. Vortex-induced dispersal of a plant pathogen by raindrop impact. Proceedings of the National Academy of Sciences 116:4917–4922.

49. Pérez-Lavalle L, Carrasco E, Vallesquino-Laguna P, Cejudo-Gómez M, Posada-Izquierdo GD, Valero A. 2021. Internalization capacity of Salmonella enterica sv Thompson in strawberry plants via root. Food Control 126:108080.

50. Reyes Esteves RG, Gerba CP, Slack DC. 2022. Control of Viral and Bacterial Contamination of Lettuce by Subsurface Drip Irrigation. Journal of Irrigation and Drainage Engineering 148.

51. Beattie GA. 2011. Water Relations in the Interaction of Foliar Bacterial Pathogens with Plants. Annu Rev Phytopathol 49:533–555.

52. Yunis H, Bashan Y, Okon Y, Henis Y. 1980. Weather Dependence, Yield Losses, and Control of Bacterial Speck of Tomato Caused by Pseudomonas tomato. Plant Dis 64:937– 939.

53. Toussaint V. 2001. Ecology of Xanthomonas campestris pv. vitians in relation to development of bacterial leaf spot of lettuce. McGill University, Montreal.

54. Panchal S, Chitrakar R, Thompson BK, Obulareddy N, Roy D, Hambright WS, Melotto M. 2016. Regulation of stomatal defense by air relative humidity. Plant Physiol 172:2021– 2032.

55. Stine SW, Song I, Choi CY, Gerba CP. 2005. Effect of relative humidity on preharvest survival of bacterial and viral pathogens on the surface of cantaloupe, lettuce, and bell peppers. J Food Prot 68:1352–1358.

56. Roy D, Panchal S, Rosa BA, Melotto M. 2013. Escherichia coli O157:H7 Induces Stronger Plant Immunity than Salmonella enterica Typhimurium SL1344. Phytopathology 103:326–332.

57. López-Gálvez F, Gil MI, Allende A. 2018. Impact of relative humidity, inoculum carrier and size, and native microbiota on Salmonella ser. Typhimurium survival in baby lettuce. Food Microbiol 70:155–161.

58. Deblais L, Helmy YA, Testen A, Vrisman C, Madrid MJ, Kathayat D. 2019. Specific Environmental Temperature and Relative Humidity Conditions and Grafting Affect the Persistence and Dissemination of Salmonella enterica subsp. enterica Serotype Typhimurium in Tomato Plant Tissues. Appl Environ Microbiol 85:1–17.

59. Walters SP, Thebo AL, Boehm AB. 2011. Impact of urbanization and agriculture on the occurrence of bacterial pathogens and stx genes in coastal waterbodies of central California. Water Res 45:1752–1762.

60. Harris CS, Tertuliano M, Rajeev S, Vellidis G, Levy K. 2018. Impact of storm runoff on *Salmonella* and *Escherichia coli* prevalence in irrigation ponds of fresh produce farms in southern Georgia. J Appl Microbiol 124:910–921.

61. Melotto M, Underwood W, He SY. 2008. Role of Stomata in Plant Innate Immunity and Foliar Bacterial Diseases. Annu Rev Phytopathol 46:101–122.

62. Kroupitski Y, Golberg D, Belausov E, Pinto R, Swartzberg D, Granot D, Sela S. 2009. Internalization of Salmonella enterica in Leaves Is Induced by Light and Involves Chemotaxis and Penetration through Open Stomata. Appl Environ Microbiol 75:6076– 6086.

63. Chincinska IA. 2021. Leaf infiltration in plant science: old method, new possibilities. Plant Methods 17:83.

64. Rufian J, Sanchez-Romero M, Lopez-Marquez D, Macho A, Mansfield J, Arnold D, Ruiz-Albert J, Casadesus J, Beuzon C. 2016. Pseudomonas syringae differentiates into phenotypically distinct subpopulations during colonization of a plant host. Environ Microbiol 18:3593–3605.

65. Newman M-A, Sundelin T, Nielsen JT, Erbs G. 2013. MAMP (microbe-associated molecular pattern) triggered immunity in plants. Front Plant Sci 4.

66. Qi J, Wang J, Gong Z, Zhou J-M. 2017. Apoplastic ROS signaling in plant immunity. Curr Opin Plant Biol 38:92–100.

67. Khan M, Ali S, Al Azzawi TNI, Saqib S, Ullah F, Ayaz A, Zaman W. 2023. The Key Roles of ROS and RNS as a Signaling Molecule in Plant–Microbe Interactions. Antioxidants 12:268.

68. Ferelli AMC, Bolten S, Szczesny B, Micallef SA. 2020. Salmonella enterica Elicits and Is Restricted by Nitric Oxide and Reactive Oxygen Species on Tomato. Front Microbiol 11:1–15.

69. García A V., Hirt H. 2014. Salmonella enterica induces and subverts the plant immune system. Front Microbiol 5:1–6.

70. Potnis N, Soto-Arias JP, Cowles KN, van Bruggen AHC, Jones JB, Barak JD. 2014. Xanthomonas perforans Colonization Influences Salmonella enterica in the Tomato Phyllosphere. Appl Environ Microbiol 80:3173–3180.

71. Steeb B, Claudi B, Burton NA, Tienz P, Schmidt A, Farhan H, Mazé A, Bumann D. 2013. Parallel Exploitation of Diverse Host Nutrients Enhances Salmonella Virulence. PLoS Pathog 9.

72. Teplitski M, Barak JD, Schneider KR. 2009. Human enteric pathogens in produce: un-answered ecological questions with direct implications for food safety. Curr Opin Biotechnol 20:166–171.

73. Abbott DW, Boraston AB. 2008. Structural Biology of Pectin Degradation by Enterobacteriaceae. Microbiology and Molecular Biology Reviews 72:301–316.

74. Chalupowicz L, Nissan G, Brandl MT, McClelland M, Sessa G, Popov G, Barash I, Manulis-Sasson S. 2018. Assessing the Ability of *Salmonella enterica* to Translocate Type III Effectors Into Plant Cells. Molecular Plant-Microbe Interactions® 31:233–239.

75. Fatima U, Senthil-Kumar M. 2015. Plant and pathogen nutrient acquisition strategies. Front Plant Sci 6:1–12.

76. Reid SJ, Abratt VR. 2005. Sucrose utilisation in bacteria: Genetic organisation and regulation. Appl Microbiol Biotechnol 67:312–321.

77. Zheng J, Allard S, Reynolds S, Millner P, Arce G, Blodgett RJ, Brown EW. 2013. Colonization and Internalization of Salmonella enterica in Tomato Plants. Appl Environ Microbiol 79:2494–2502.

78. Smith NW, Shorten PR, Altermann E, Roy NC, McNabb WC. 2019. The Classification and Evolution of Bacterial Cross-Feeding. Front Ecol Evol 7.

79. Murillo-Roos M, Abdullah HSM, Debbar M, Ueberschaar N, Agler MT. 2022. Cross-feeding niches among commensal leaf bacteria are shaped by the interaction of strain-level diversity and resource availability. ISME J 16:2280–2289.

80. Palaniraj A, Jayaraman V. 2011. Production, recovery and applications of xanthan gum by Xanthomonas campestris. J Food Eng 106:1–12.

81. Lovelace AH, Ma W. 2022. How do bacteria transform plants into their oasis? Cell Host Microbe 30:412–414.

82. Cowles KN, Willis DK, Engel TN, Jones JB, Barak JD. 2016. Diguanylate cyclases AdrA and STM1987 regulate salmonella enterica exopolysaccharide production during plant colonization in an environment-dependent manner. Appl Environ Microbiol 82:1237– 1248.

83. Barak J, Whitehand L, Charkowski A. 2002. Differences in Attachment of Salmonella enterica Serovars and Escherichia coli O157:H7 to Alfalfa Sprouts. Appl Environ Microbiol 68:4758 LP – 4763.

