## Supplemental Fig 1-6 for "Time of arrival during plant disease progression and humidity additively influence *Salmonella enterica* colonization of lettuce"

**SUPPLEMENTARY MATERIALS**

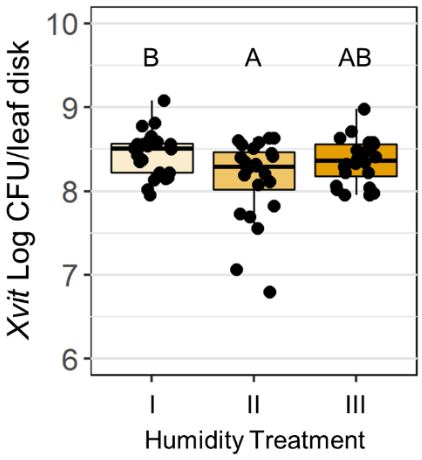


**Supplementary Figure 1**: Humidity fluctuation mildly impacts *X. vitians* populations *in planta*. *X. vitians* populations *in planta*. *X. vitians*-infiltrated lettuce plants were divided into three treatments groups, as described in **Figure 2**: I: 50-70% RH for 7 days; II: 50-70% RH for 4 days, >90% RH for 1 day, 50-70% RH for 2 days; III: 50-70% RH for 4 days, >90% RH for 3 days. Boxplots show *X. vitians* populations at 7 days post-infiltration. Letters above the boxplots indicate significance between treatment groups based on Welch-ANOVA and *post hoc* Games-Howell means comparison tests ($P < 0.05$). $N = 24$ leaves/treatment group; data is pooled from three independent experimental replicates.

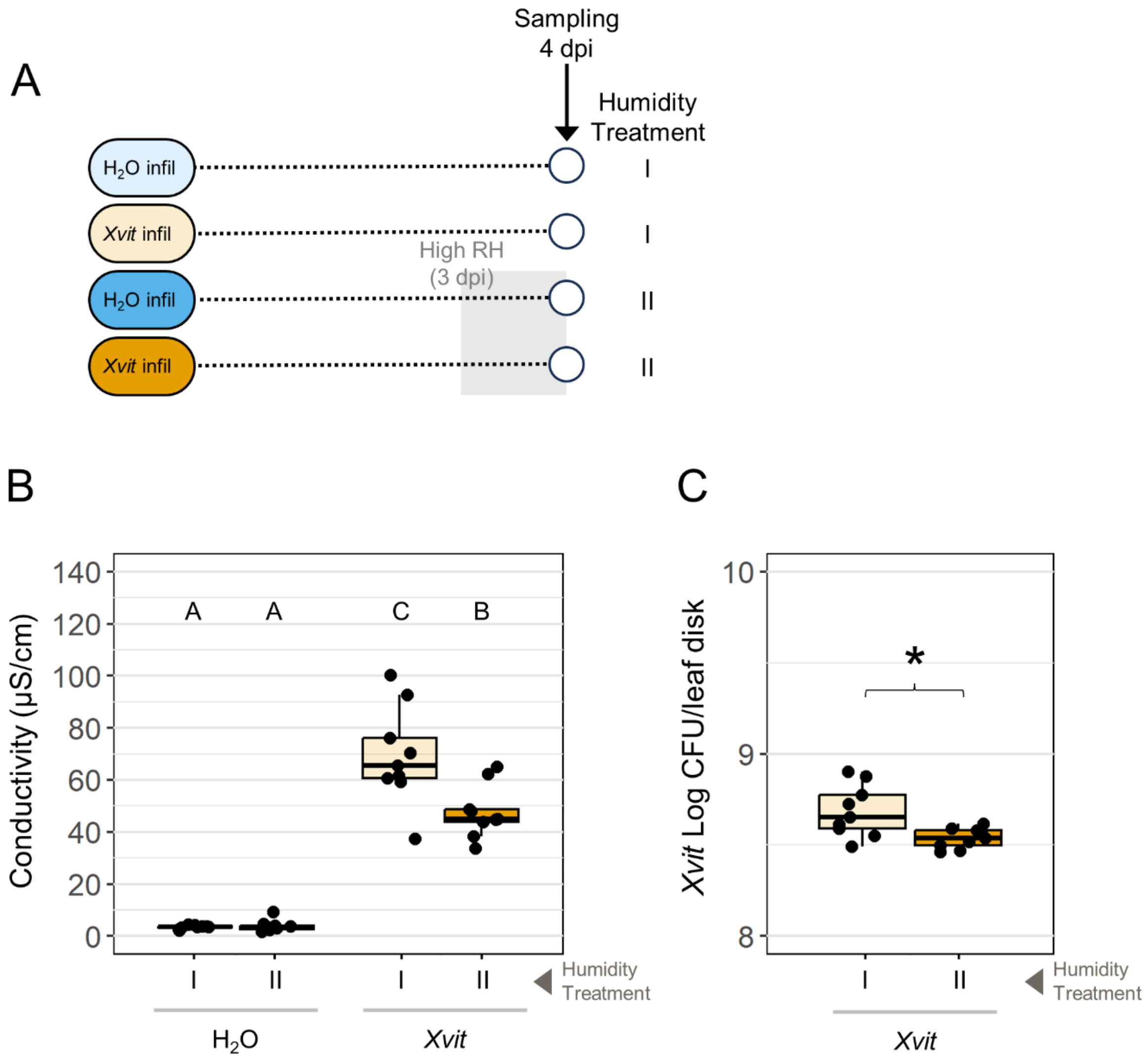


**Supplementary Figure 2:** The introduction of high humidity during early infection impacts disease caused by *X. vitians*. **(A)** Schematic of experimental design. Plants were sampled at 4 days post-infiltration with $H_2O$ or *X. vitians* cells after 0 or 24 hours of exposure to high humidity (>90%). **(B)** Electrolyte leakage and **(C)** *X. vitians in planta* colonization of lettuce leaves. Letters above the boxplots indicate significance between treatment groups based on Welch-ANOVA and *post hoc* Games-Howell means comparison tests, and asterisk represents significant difference based on a Welch T-test ($P < 0.05$). $N = 9$ leaves/treatment group; data is pooled from three independent experimental replicates.

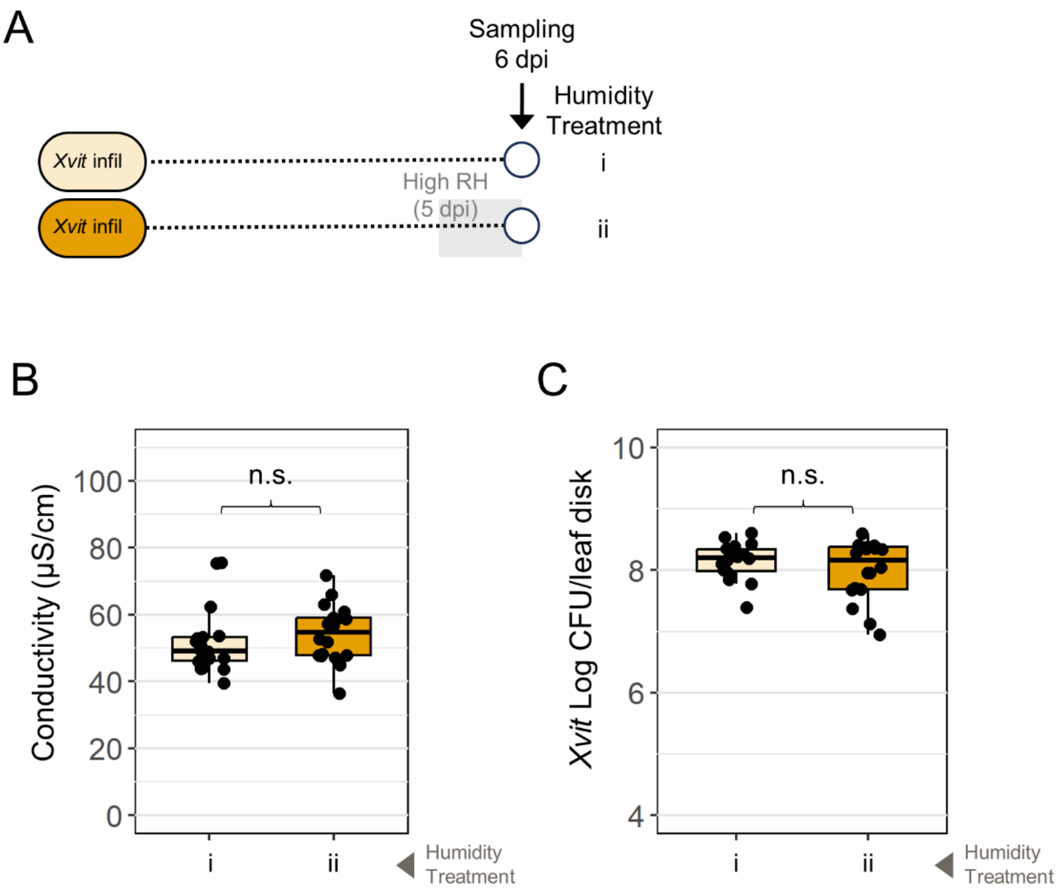


**Supplementary Figure 3:** The introduction of high humidity during late infection does not impact disease caused by *X. vitians*. **(A)** Schematic of experimental design. Plants were sampled at 6 days post-infiltration (dpi) with *X. vitians* cells after 0 or 24 hours of exposure to high humidity (>90%) applied at 5 dpi. **(B)** Electrolyte leakage and **(C)** *X. vitians in planta* colonization of lettuce leaves. Significance testing based on a Welch T-test ($P < 0.05$); n.s. = not significant. $N$ = 18 leaves/treatment group; data is pooled from three independent experimental replicates.

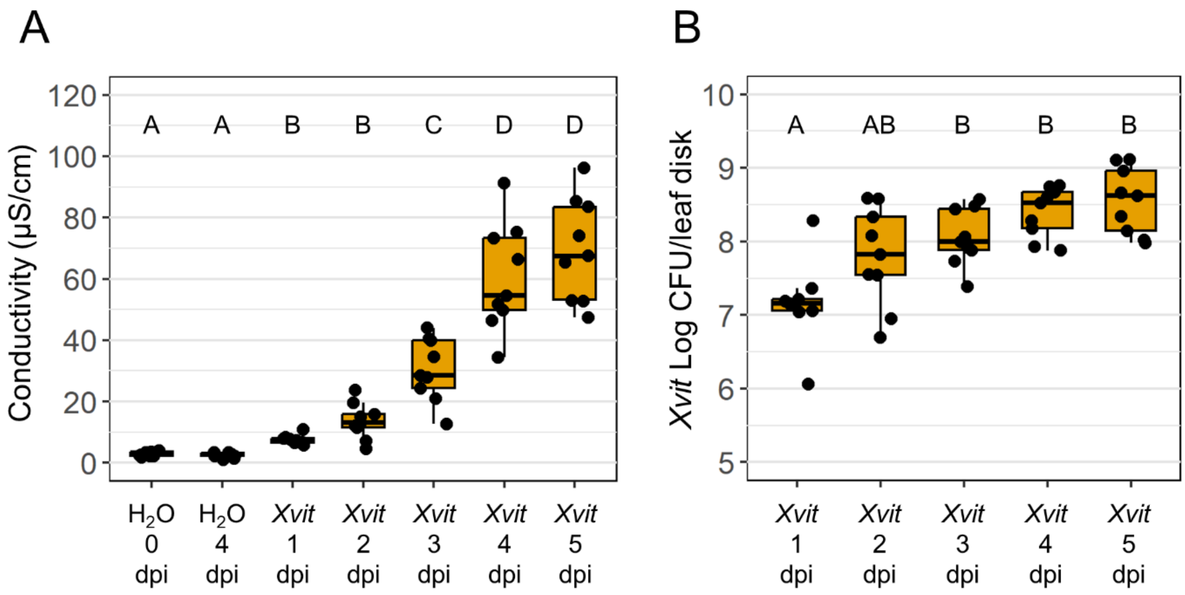


**Supplementary Figure 4:** *X. vitians* infection induces electrolyte leakage and supports *X. vitians* replication in the lettuce host at moderate humidity. **(A)** Electrolyte leakage and **(B)** *X. vitians in planta* colonization of lettuce leaves at the given days post-infiltration (dpi) and for the given infiltration treatment ($H_2O$ or *X. vitians*). Plants were maintained at a humidity of 50-70% for the duration of the experiment. Letters above the boxplots indicate significance between treatment groups based on Welch-ANOVA and *post hoc* Games-Howell means comparison tests ($P < 0.05$). $N = 9$ leaves/treatment group; data is pooled from three independent experimental replicates.

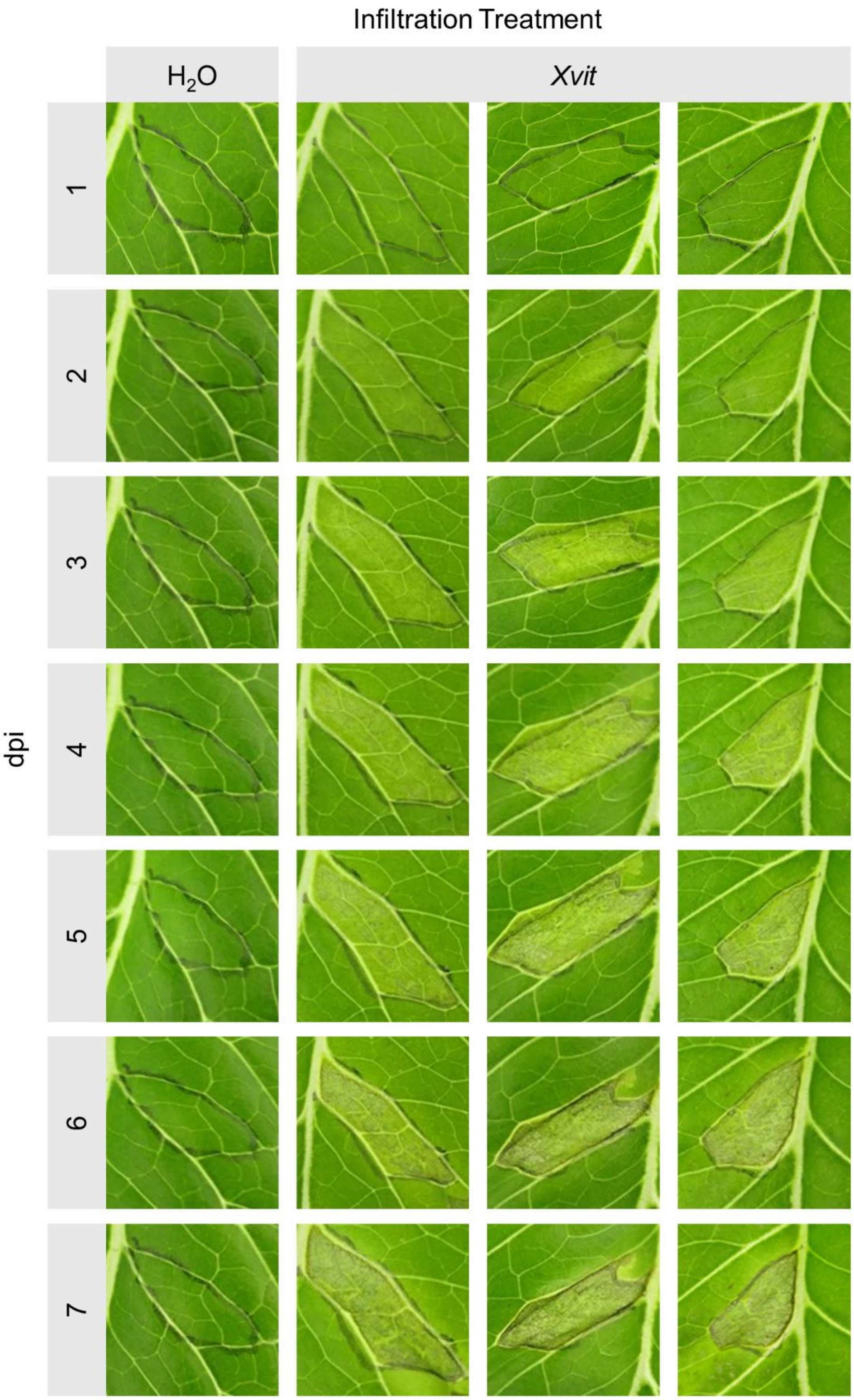
Infiltration Treatment
H2O
Xvit
dpi
1
2
3
4
5
6
7

**Supplementary Figure 5 (previous page):** Infiltration with *X. vitians* results in progression of water soaking and necrosis. Daily time course images are shown for one representative $H_2O$-infiltrated plant and three representative *X. vitians* (*Xvit*)-infiltrated lettuce plants at 1-7 days post-infiltration (dpi). Relative humidity was maintained at 50-70%. *Xvit*-infiltrated plants were infiltrated with a *X. vitians* culture prepared to $3x10^8$ CFU/mL.

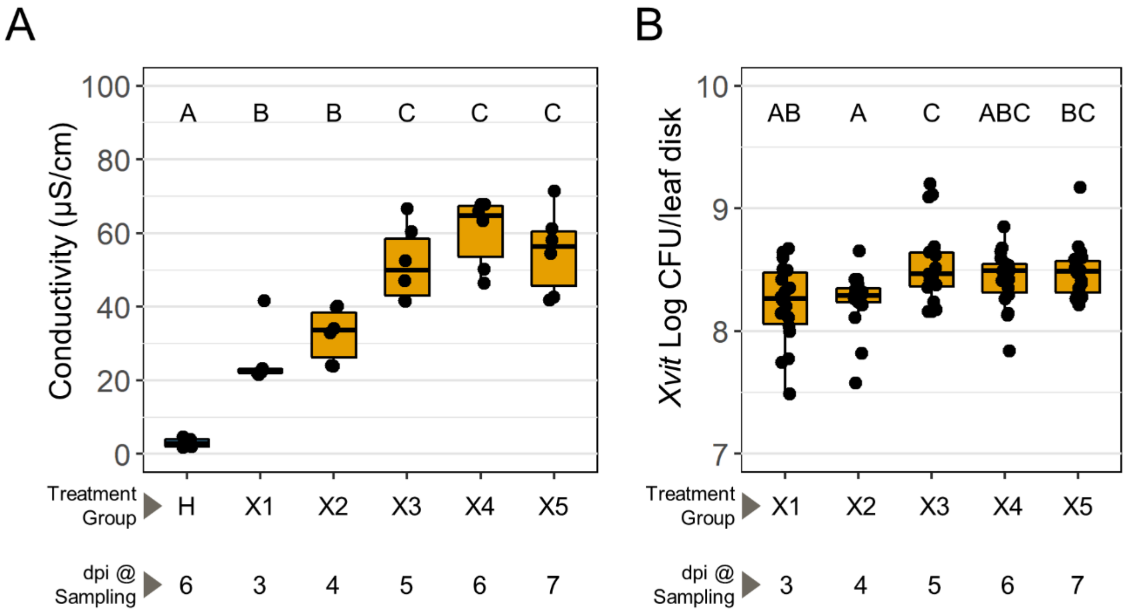


**Supplementary Figure 6:** *X. vitians* infection induces electrolyte leakage in lettuce host. **(A)** Electrolyte leakage and **(B)** *X. vitians in planta* colonization of lettuce leaves sampled two days post-arrival of *S. enterica*, as described in **Figure 4**. Plants were exposed to high humidity (>90% RH) for the 24 hours immediately prior to sampling. Data for *X. vitians*-infiltrated lettuce represents infection state at 3-7 days post-infiltration (dpi) (Treatments X1-X5). $H_2O$-infiltrated plants were sampled at 6 dpi. Letters above the boxplots indicate significance between treatment groups based on Welch-ANOVA and *post hoc* Games-Howell means comparison tests ($P < 0.05$). For (A), $N$ = 5-6 leaves/treatment group; data represents one representative experimental replicate. For (B), $N$ = 18 leaves/treatment group; data is pooled from three independent experimental replicates.

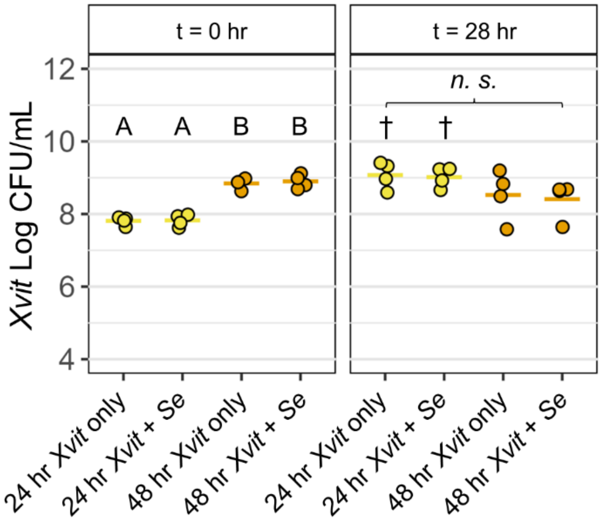


**Supplementary Figure 7:** S. enterica does not impact X. vitians cell concentrations *in vitro*. *X. vitians* cultures grown for 24 (yellow) or 48 hours (orange) were mixed with water or *S. enterica* cells at time (t) = 0 hr. Each horizontal bar represents the average *X. vitians* cell concentrations of four independent experiments. Letters above the boxplots or n.s. (not significant) indicate significance between treatment groups based on Welch-ANOVA and *post hoc* Games-Howell means comparison tests ($P < 0.05$) within each time point. Daggers (†) indicate a significant increase in *X. vitians* cell concentration at time = 28 hr relative to time = 0 hr within each treatment based on two-sample T-tests ($P < 0.05$).
